# Reconstituting centriole biogenesis on an engineered cellular platform uncovers organelle assembly principles

**DOI:** 10.64898/2026.08.22.746443

**Authors:** Samya Aich, Pierre Gönczy

**Affiliations:** Swiss Institute for Experimental Cancer Research (ISREC), School of Life Sciences, Swiss Federal Institute of Technology Lausanne (EPFL), Lausanne, Switzerland

**Keywords:** Centriole assembly, organelle biogenesis, number control, torus, phase-separation, synthetic droplet, expansion microscopy

## Abstract

Centriole copy number is tightly regulated, with one procentriole emanating from a torus surrounding each pre-existing centriole. Which proteins are sufficient to generate a full-fledged organelle is unclear. We address this question by engineering a high valency low copy number phase-separated droplet platform to concentrate proteins in an ectopic cellular location. We establish that droplet targeting of the torus protein Cep63 suffices to initiate procentriole assembly. Ectopic procentrioles mature when a limiting interaction involving STIL is alleviated or when pre-existing centrioles are lacking. Ectopic procentrioles disengage from the droplet during mitosis in a PLK1-dependent manner, organize supernumerary spindle poles, and seed procentriole formation at the next cell cycle, demonstrating that synthetic centrioles have been reconstituted. Moreover, we uncover that procentriole number scales with platform surface area. Since the torus surface area is set by pre-existing centriole dimensions, we propose that this constitutes a closed circuit mechanism dictating organelle number homeostasis.

**Highlights:**

□ A high valency low copy number droplet platform engineered for cellular reconstitution
□ Targeting the torus protein Cep63 to the droplet leads to naked cartwheel assembly
□ Eliminating competition from resident centrioles enables synthetic centriole reconstitution
□ Synthetic centriole number scales linearly with reconstituted torus surface area

## Introduction

Centrioles are evolutionarily conserved cylindrical macromolecular assemblies containing over one hundred protein types organized in a precise architecture, including a signature nine-fold radial symmetry of microtubules. Together with the surrounding peri-centriolar material (PCM), centrioles constitute the centrosome, an important microtubule organizing center (MTOC) of animal cells, and thereby contribute to microtubule-based processes, including bipolar spindle formation during mitosis^1,2^. Just like the genetic material, centrioles are subject to tight copy number control in cycling cells^1,2^. G1 cells possess two pre-existing centrioles (resident centrioles hereafter), which each seed the formation of one procentriole in their vicinity towards the G1/S transition, such that mitotic cells possess four centriolar cylinders, two per spindle pole. Supernumerary centrioles can result in multipolar spindle assembly and fuel genome instability, underscoring the importance of centriole number homeostasis.

A critical control point occurs at the onset of procentriole assembly, when a cartwheel element emanates from a protein complex termed the torus, which surrounds the proximal part of each resident centriole. The torus is organized in a stereotyped arrangement of four interacting proteins, with Cep57 and Cep57L1 closest to the resident centriole, followed by Cep63, and then the outermost protein Cep152^3–9^. Cep152 is thought to recruit the kinase PLK4^10–13^, which together with its substrate STIL, focuses on the torus surface^14–17^. The PLK4/STIL focus promotes formation of a 9-fold symmetric cartwheel comprising phosphorylated STIL and its interacting partner, the cartwheel building block protein SAS-6^18–21^. Phosphorylated STIL also recruits the microtubule-binding protein CPAP, which is critical for processive elongation of centriolar microtubules^22–27^. Each procentriole remains juxtaposed next to the resident centriole until mitosis, when the two disengage in a PLK1-dependent manner^28–31^. PLK1 also ensures that the newly made procentriole acquires the centriolar protein Cep295^32–34^, which is essential for conversion of the procentriole into a full-fledged centriole capable of seeding a new procentriole in turn at the next cell cycle^1,35,36^.

What mechanisms operate in cycling cells to ensure that just one procentriole forms per resident centriole, and strictly at that location? These mechanisms include limiting the levels of critical components that act at the onset of procentriole formation. Indeed, overexpression of PLK4, STIL or SAS-6 results in supernumerary centrioles, which form invariably around the torus of resident centrioles, indicating that increased protein levels do not suffice normally to trigger organelle formation elsewhere in the cell^15,37,38^. By contrast, in human cells experimentally depleted of resident centrioles, several procentrioles can form *de novo* in the cytoplasm^39,40^, indicating that organelle biogenesis can occur elsewhere than on the torus under some circumstances. Interestingly, the presence of just one resident centriole prevents such *de novo* formation, with a procentriole emanating from the torus of the single resident centriole in this case^40^. An attractive hypothesis to explain such spatial restriction is that the cytoplasmic levels of critical proteins triggering procentriole assembly are normally sufficiently low to prevent illegitimate formation elsewhere, with the torus acting to concentrate such proteins^41^. Echoing this possibility, in differentiating multiciliated cells, several procentrioles emanate from the deuterosome, a large electron-dense spherical structure that is proposed to stem from the younger resident centriole and comprises notably the Cep63 paralogue Deup1^42,43^. These post-mitotic cells have a distinct cell cycle state and undergo global transcriptional upregulation of centriolar proteins, including PLK4, STIL and SAS-6^44^, thereby likely contributing to the ability of the deuterosome to yield several procentrioles.

In cycling cells, by contrast, it is unclear whether torus proteins instruct centriole assembly and contribute to number control. This paucity of knowledge reflects notably the fact that the contribution of individual proteins is difficult to ascertain because of redundancies. For example, despite the likely pivotal role of the four torus proteins, none of them is unequivocally essential for centriole biogenesis^5,10,11,45–47^. We reasoned that a test of sufficiency, in which individual proteins are rerouted to an ectopic cellular location to reconstitute centriole assembly, would prove instructive in unraveling the contribution of individual torus proteins, as well as in uncovering general principles of organelle biogenesis.

## Results

### Engineering an ectopic torus surface in the cellular context

We sought to engineer a modular platform to which individual proteins can be targeted to test their sufficiency in cellular assembly reactions, including centriole biogenesis. We reasoned that such a platform must satisfy two critical requirements. First, to achieve high local concentration of targeted proteins, a high valency platform with densely packed binding sites is needed. Second, to avoid diluting endogenous interactors between several sites, a small and low copy number platform is needed. Moreover, the platform must not interfere with cellular function. With these goals in mind, we functionalized the synthetic phase-separating construct Sumo_(10)_-SIM_(6)48_ with GFP, thereby engineering a cellular assembly platform enabling targeting proteins of interest tagged with GFP Binding Protein (GBP) to the surface of Sumo_(10)_-SIM_(6)_-GFP phase-separated droplets (Figure 1A). Due to homotypic fusion, these droplets are present in one or two copies in most cells, with a mean diameter of ∼1.5 μm (Figures S1A, S1B). These droplets persist through mitosis and do not interfere with cell-cycle progression (Figures S1C, S1D), together making them an ideal high valency low copy number platform to ectopically concentrate proteins of interest. To streamline the usage of this platform, we generated stable U-2 OS FlpIn cells harboring an integrated polycistronic vector constitutively expressing GFP-Sumo_(10)_-SIM_(6)_-GFP, as well as doxycycline inducible GBP-tagged proteins (Methods).

**Figure 1.**
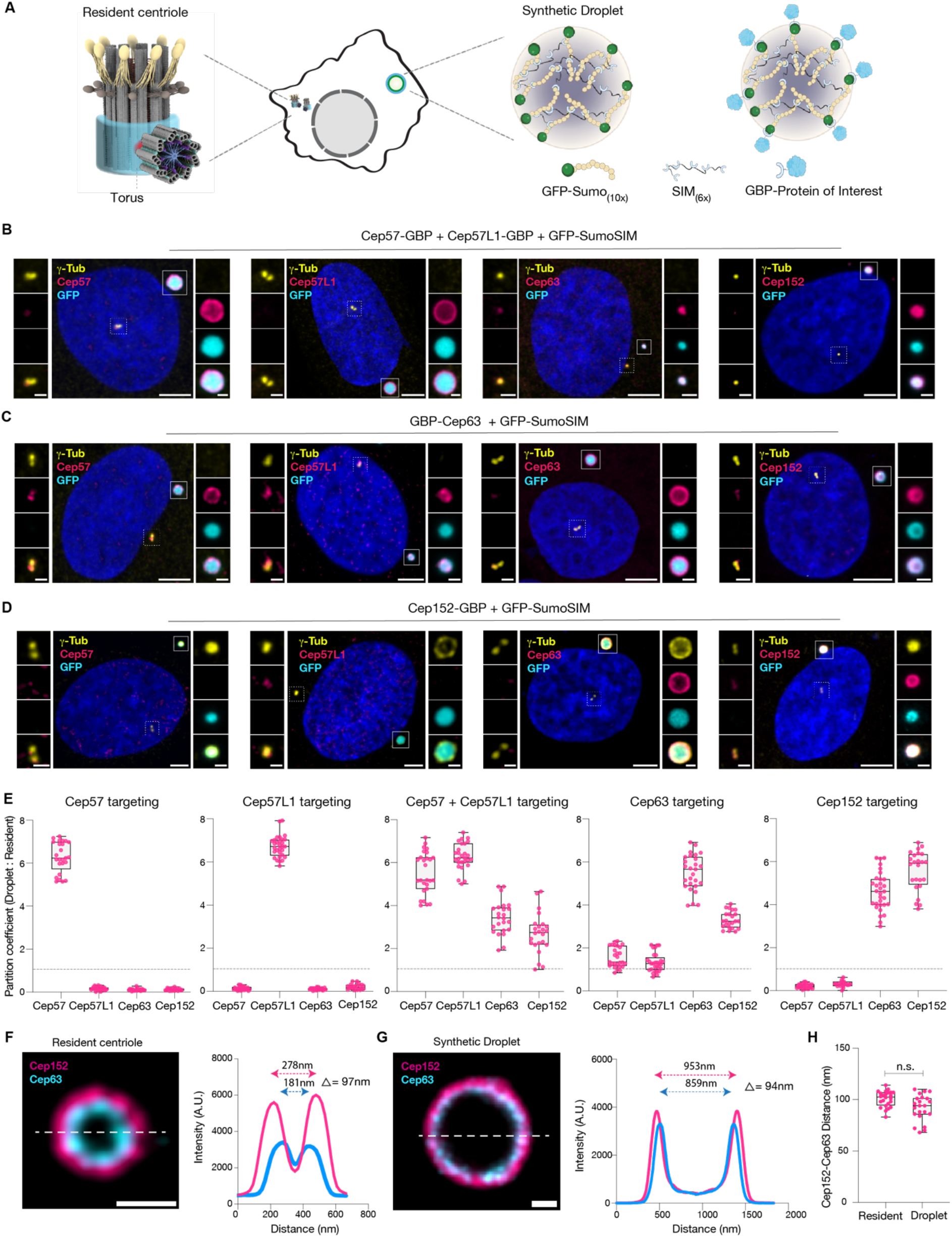
Interdependence of torus proteins upon droplet targeting. **A**) Schematic of droplet-based rerouting strategy for engineered centriole assembly. Left: resident centriole with surrounding torus (blue). Right: synthetic droplet comprising tandem repeats of GFP-Sumo_(10x)_ and SIM_(6x)_ that recruits GBP (GFP binding protein)-tagged Protein of Interest (POI) to its surface. **B**-**D**) Representative immunofluorescence images of cells with droplet targeted with indicated GBP tagged torus protein and stained with antibodies against indicated proteins. γ-tubulin marks resident centrioles, GFP the droplet. Here and in all other figures, left insets show magnified views of resident centrioles (white dashed box), right insets magnified view of the droplet (solid white box). Note that Cep152 recruits γ-tubulin to the droplet, revealing a hitherto unsuspected sufficiency in localizing this PCM protein. **E**) Partition coefficient of torus proteins upon droplet-targeting. Partition coefficient: mean intensity of each torus protein at the droplet normalized to mean intensity at resident centrioles in the same cell. Each data point represents one cell. Boxes, horizontal lines and whiskers represent interquartile ranges, median and minima-maxima. Dashed line indicates partition coefficient of 1. At least 25 cells for each condition were pooled from three independent repeats. **F, G**) Representative single section U-ExM images of resident centriole (F) and droplet (G), immunostained for Cep63 and Cep152. Dashed line indicates position of the line used to quantify diameter as peak-to-peak distance. Resident centriole is from a neighboring cell not containing a droplet, because the depletion of Cep152 at the resident centriole in cells with droplet results in a signal that is too low for robust size measurements. **H**) Difference between diameters of Cep63 and Cep152 protein distributions. n=22 (resident centriole) and n=24 (droplet), pooled from two independent experiments. Only top views were measured for resident centrioles. Two-tailed t-test was used to calculate statistical significance. n.s. indicates p >0.05. Scale bar in B-D: 5 μm in low magnification views, 1 μm in insets. Scale bar in F, G: 250 nm.

We first used this platform to systematically investigate the ability of GBP-tagged proteins of the torus to recruit one another (Figure S2A). To distinguish ectopic assemblies forming at the droplet from procentriole forming next to resident centrioles in the same cell, we immunostained the PCM marker γ-tubulin to mark resident centrioles, and GFP to label the droplet. Using this approach, we found that targeting either Cep57 or Cep57L1 to the droplet is not sufficient to recruit Cep63 or Cep152 (Figure 1E; Figures S2B, S2C). By contrast, when Cep57 and Cep57L1 are targeted jointly to the droplet, Cep63 and Cep152 are robustly recruited as well (Figures 1B, 1E). Importantly, when Cep63 is targeted to the droplet, it alone recruits Cep57, Cep57L1 and Cep152 (Figures 1C, 1E). By contrast, although droplet-targeted Cep152 can recruit Cep63, neither Cep57 nor Cep57L1 is detected in this case (Figures 1D, 1E). Overall, upon targeting to a high valency low copy number synthetic droplet, Cep57 and Cep57L1 jointly, as well as Cep63 alone, can reconstitute an ectopic surface with a full complement of torus proteins.

Since Cep63 alone is sufficient to recruit all other torus proteins, we prioritized it to investigate whether it recapitulates features of the endogenous torus surface. Cep152 is positioned peripheral to Cep63 on the torus surrounding resident centrioles^4,6^. We set out to address whether this arrangement is preserved on the droplet using ultrastructure expansion microscopy (U-ExM) for increased resolution and found this to be the case indeed (Figures 1F-H). Additionally, we observed that upon GBP-Cep63 targeting to the droplet, Cep152 is depleted from resident centrioles, albeit without impacting procentriole formation (Figures S2D-S2F). Together, these findings indicate that Cep63 targeting to the droplet reconstitutes an ectopic torus-like surface, without affecting centriole biogenesis from resident centrioles.

### Cep63 triggers cartwheel assembly

We next tested whether such an ectopic torus-like surface is sufficient to trigger procentriole assembly. Remarkably, we found that targeting Cep63 to the droplet is sufficient to recruit PLK4 in ∼80% of cells in which PLK4 is also present at resident centrioles (Figures 2A-C). Moreover, the PLK4 substrate STIL is likewise recruited to the droplet, as is the cartwheel protein SAS-6 (Figures 2A-C). Importantly, rather than being distributed uniformly on the droplet, PLK4, STIL and SAS-6 localize to one or a few foci (Figure 2A), indicative of organized assemblies rather than merely passive protein accumulation. These findings establish that concentrating Cep63 is sufficient to efficiently recruit PLK4, STIL and SAS-6 to foci on the droplet surface.

**Figure 2.**
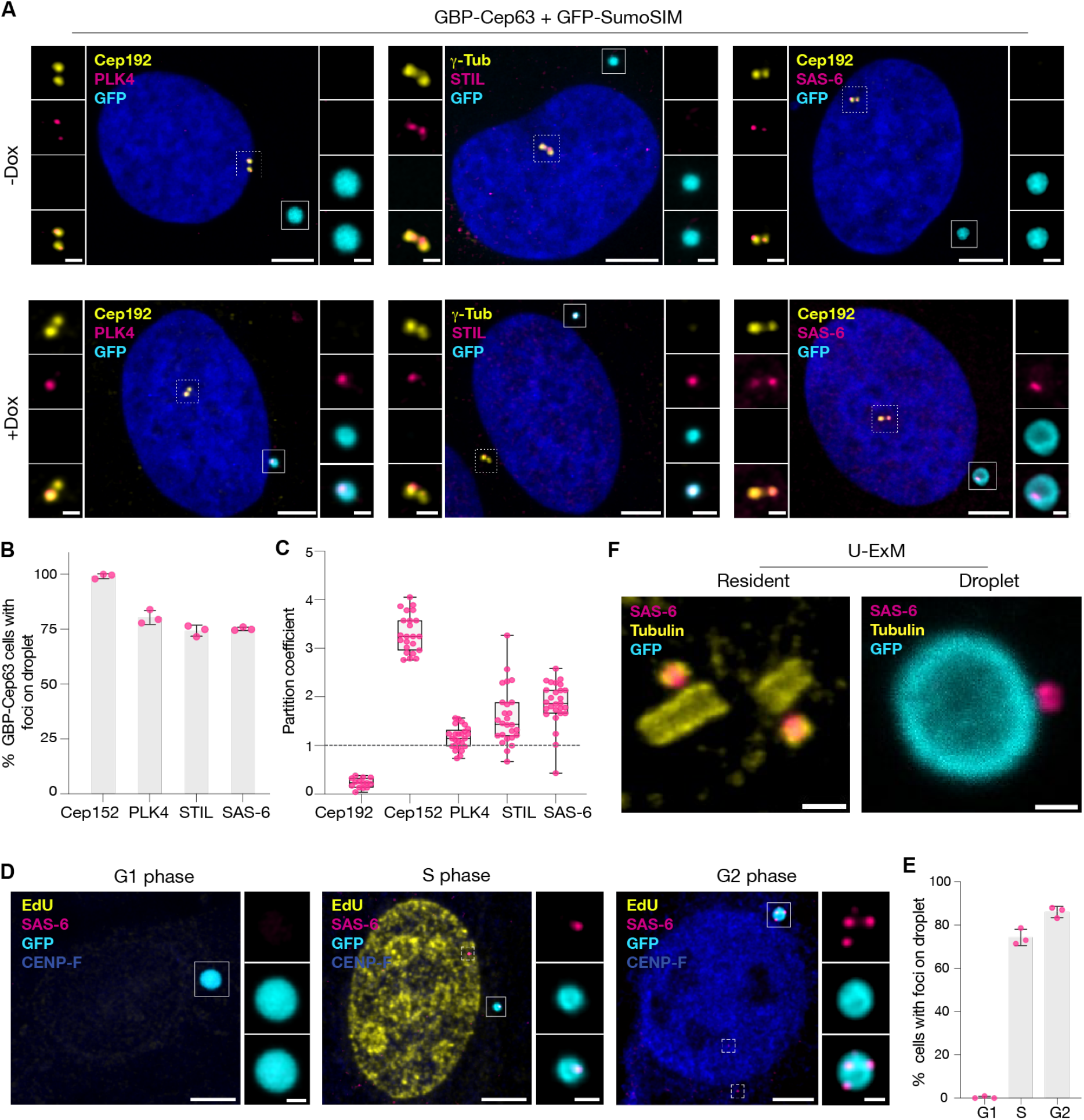
Targeting Cep63 to the droplet is sufficient for cartwheel assembly. **A)** Representative images of droplet-containing interphase U-2 OS cells without (-Dox, top) or with (+Dox, bottom) doxycycline-mediated GBP-Cep63 expression, immunostained for indicated centriolar proteins. Cep192 or γ-tubulin was used to mark resident centrioles. **B)** Percentage of droplet targeted Cep63 cells with droplets harboring indicated centriolar proteins as a fraction of cells where the same centriolar proteins are present at resident centrioles. At least 100 cells were scored for each condition per experiment in three independent experiments. Bars represent mean +/- s.d. **C)** Partition coefficient of indicated centriolar proteins between droplet and resident centrioles. Each data point represents an individual cell. For droplets with multiple foci, an average mean intensity is used. Boxes represent interquartile ranges, horizontal line median, whiskers minima and maxima. n=14 (Cep192), 23 (Cep152), 25 (PLK4), 27 (SAS-6) and 26 (STIL), pooled from three independent experiments. Note that Cep192 is not recruited to the droplet, exemplifying specificity in the recruitment of the other proteins. **D)** Cells immunostained for SAS-6 together with the S phase marker EdU and the G2 phase marker CENP-F. **E)** Percentage of cells with SAS-6 foci on droplet at indicated cell cycle phases. 50 cells were counted for each condition per experiment in three independent experiments. Bars represent mean +/- s.d. **F)** Representative max-projected U-ExM images of resident centrioles and droplet from same cell, immunostained for α/β-tubulin, GFP and SAS-6. Note absence of tubulin on the droplet. Scale bars in A and D: 5 μm in low magnification views, 1 μm in insets. Scale bars in F: 250 nm.

Procentriole formation at resident centrioles initiates towards the G1/S transition, such that the SAS-6 based cartwheel is absent from most cells in G1, but present in S and G2^49^. We asked whether the SAS-6 foci forming on the droplet are also present in such a cell-cycle dependent manner. Co-staining with the S phase marker EdU and the G2 marker CENP-F demonstrated that this is the case, with SAS-6 foci being absent during G1 (EdU and CENP-F negative), but present in the vast majority of cells in S (EdU positive and CENP-F negative) and G2 (EdU negative and CENP-F positive) (Figures 2D, 2E).

We also asked whether foci forming on the droplet exhibit dimensions of assemblies present during physiological procentriole assembly. Strikingly, analysis by U-ExM revealed that SAS-6 foci on the droplet surface exhibit indistinguishable dimensions from procentrioles formed next to resident centrioles in the same cell (Figures 2F; Figure S3A). Similar results were obtained for STIL (Figure S3B). Strikingly, however, we found that these cartwheel-like elements are invariably devoid of tubulin (Figure 2F). Accordingly, the centriolar microtubule-interacting proteins CPAP, CP110 and Cep295 are also absent from droplet foci in most cells (Figures S3C, S3D).

Together, these findings demonstrate that the torus-like surface organized by droplet-targeted Cep63 is sufficient to generate ectopic foci of PLK4 and STIL, as well as cartwheels containing STIL and SAS-6. However, these assemblies are devoid of more distal and peripheral components of maturing procentrioles, including microtubules, and likely represents the “naked cartwheel” intermediate observed in early stages of physiological procentriole formation^50^.

### Ectopic procentrioles develop further upon excess STIL

What prevents the cartwheel formed upon Cep63 droplet targeting from recruiting microtubules and extending further? We reasoned that an interaction normally occurring between a protein present on the cartwheel and a protein important for subsequent assembly steps might be limiting. Such an interaction could involve one of the microtubule-interacting proteins that we found to be lacking from the ectopic naked cartwheel. A strong such candidate is CPAP, which is normally recruited to the growing procentriole through an interaction with phosphorylated STIL^22,23,51^.

Previous work established that the interaction between STIL and CPAP is inhibited by the RNA binding protein RBM14, with RBM14 depletion resulting in ectopic cytoplasmic CPAP-containing structures that form supernumerary spindle poles during mitosis (Figures S4A-S4D)^52^. If the interaction between STIL and CPAP is limiting, then RBM14 depletion should facilitate CPAP recruitment to the assemblies present on the Cep63-targeted droplet and result in further procentriole maturation. Importantly, we found this to be the case indeed (Figures 3A, 3C). Moreover, CP110 and Cep295 are also recruited onto ectopic foci upon RBM14 depletion, suggesting further maturation of the initial assemblies (Figures 3A, 3C). We reasoned that if RBM14 functions by limiting the interaction between STIL and CPAP, then this limitation should also be alleviated by overexpressing STIL. Accordingly, we found that overexpressing SNAP-STIL likewise leads to robust recruitment of CPAP, CP110 and Cep295 to Cep63-containing droplets (Figures 3B, 3C). Note that in such conditions many more foci typically appear on the droplet, as they do at resident centrioles upon excess of STIL^15^.

**Figure 3.**
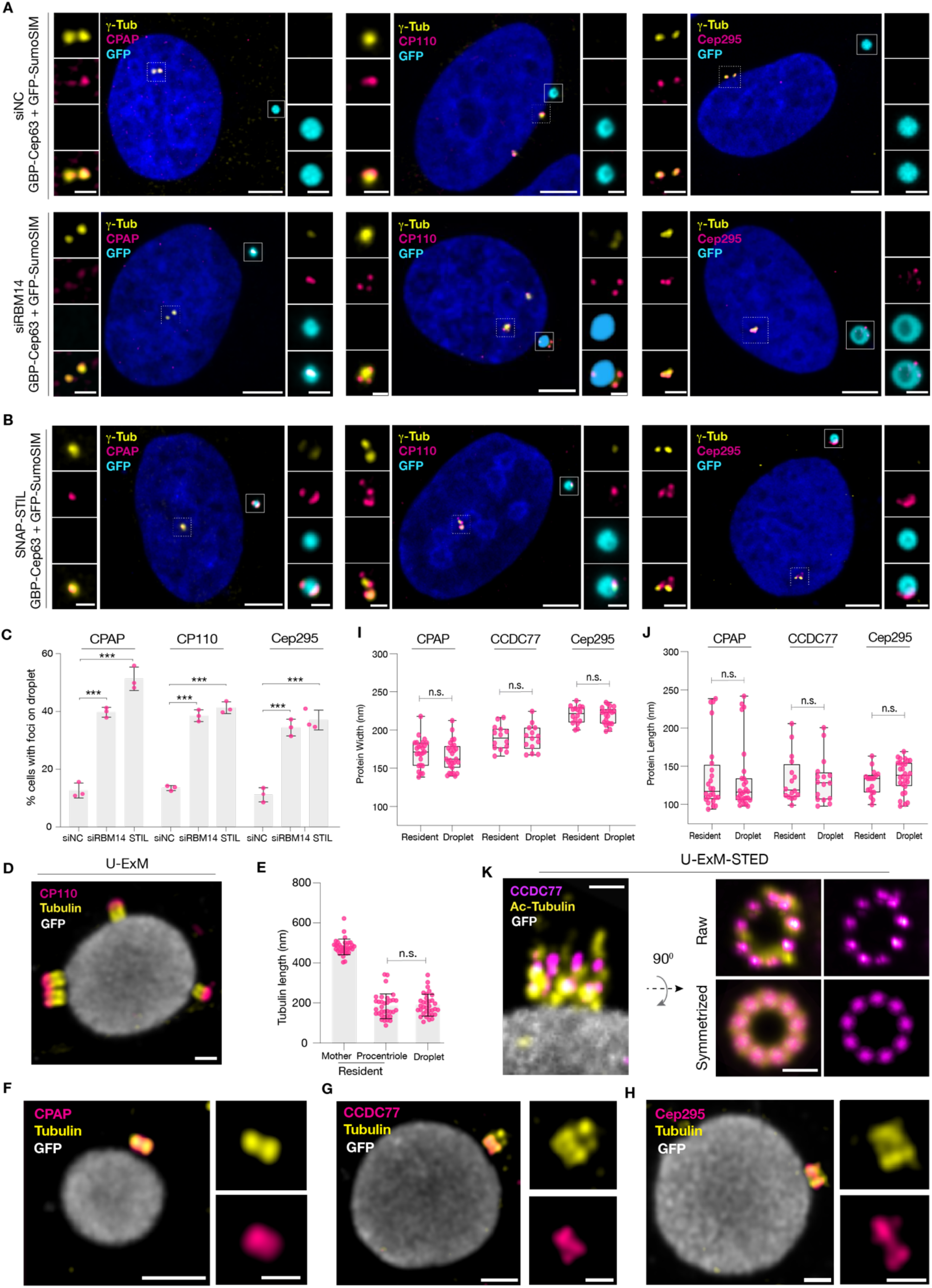
RBM14 depletion or STIL overexpression allows droplet-targeted Cep63 to form ectopic procentrioles. **A)** Representative images of cells with Cep63 targeted to the droplet and treated with either negative control siRNAs (siNC) or siRNAs against RBM14 (siRBM14), followed by immunostaining for indicated centriolar proteins. **B)** Representative images of cells with Cep63 targeted to the droplet, along with global over-expression of SNAP-STIL. **C)** Percentage of cells with droplet co-localizing with foci of indicated centriolar proteins. At least 100 cells were scored for each condition in a single experiment; three independent experiments. Bars represent mean +/- s.d. Note that in contrast to Fig. 2B, here all cells of the asynchronous population were analyzed, because frequent centriole overduplication at resident centrioles prevents reliable assessment of cell cycle state based on procentriole status at that location. Statistical significance was calculated with one-way ANOVA with Dunnett’s test for multiple comparisons. *** indicates p <0.0001. **D)** Max projected U-ExM image of droplets stained for tubulin and CP110. **E)** Length of procentriole tubulin signal measured at resident centrioles and at droplet in same cell. n= 34 cells were scored from three independent experiments. Bars represent mean +/- s.d. Paired two-tailed t-test was used to calculate statistical significance. **F-H**) Max-projected U-ExM images of droplets stained for tubulin and indicated centriolar proteins. **I, J**) Length and width of CPAP, Cep295 and CCDC77 procentriole signals at resident centrioles compared to the droplet. Each data point represents an individual cell, boxes represent interquartile ranges, horizontal line median, whiskers minima and maxima. n = 23 (CPAP), 16 (CCDC77) and 28 (Cep295), pooled from two independent experiments. Paired two-tailed t-test was used to calculate statistical significance. n.s. indicates p >0.05. **K**) Representative U-ExM-STED image of procentrioles at the droplet stained for CCDC77 (magenta) and acetylated tubulin (yellow). A volume projection on the right is shown to demonstrate the 9-fold symmetry of CCDC77. Images representative of n=12. Scale bars in A and B: 5 μm in low magnification views, 1 μm in insets. Scale bars in D, F-H: 250 nm. Scale bar in K: 100 nm.

The coordinated localization of CPAP, CP110 and Cep295 under the above conditions led us to test whether these further assemblies resemble full-fledged procentrioles. Analysis with U-ExM revealed microtubule-containing cylindrical structures growing from the surface of the droplet and capped by CP110 on their distal end (Figure 3D). Pairwise quantification of microtubule lengths between droplet-based structures and procentrioles adjacent to resident centrioles revealed similar dimensions (Figure 3D, Figure S5A). Therefore, ectopic procentrioles mature in synchrony with those next to resident centrioles.

We also used U-ExM to investigate whether proteins characteristic of mature procentrioles are likewise present in ectopic droplet-based elements. Accordingly, we found that like endogenous procentrioles^53^, droplet-based microtubules are acetylated and mono-glutamylated (Figures S5B, S5C), but lack branched chain glutamylation (Figure S5D). We also found that CPAP^54^, the A-C linker protein CCDC77^50,55^ and the microtubule-binding protein Cep295^33,34^ all localize along the proximal microtubule wall, as expected (Figures 3F-J). Furthermore, we found that the pinhead protein Cep44^56^, the putative triplet base protein Cep135^50^ and the central core component Centrin^50,57^ each occupy their expected locations (Figures S5E-S5I). To address whether the ectopic assemblies exhibit the signature 9-fold radial symmetry of centriolar cylinders, we coupled U-ExM to STED microscopy for further enhanced resolution, using the discrete spot of the A-C linker protein CCDC77 as a marker of fold symmetry. Remarkably, this analysis revealed that procentriole assemblies at the droplet exhibit 9-fold symmetry (Figure 3K).

Together, our data indicate that Cep63 is sufficient to instruct the formation of mature ectopic procentrioles with the proper architecture when STIL is present in sufficient excess to overcome inhibition by RBM14 and thereby enable CPAP recruitment.

### Ectopic procentrioles disengage from the droplet during mitosis

We set out to address whether procentrioles assembled on the droplet exhibit functional properties that characterize endogenous procentrioles, starting with an investigation of ectopic procentriole fate during mitosis. Immunostaining with CP110 revealed that most droplets in G2 cells harbor tightly associated procentrioles, whereas this is less frequent in prophase, and never the case in metaphase (Figures 4A, 4B; Figure S6A). Two possibilities can explain these observations: either ectopic procentrioles dissociate from the droplet and persist elsewhere in the cell, or they are eliminated. To distinguish between these possibilities, we conducted live imaging of U-2 OS cells stably expressing RFP-Centrin1^58^ and induced to express GBP-Cep63, SNAP-STIL and GFP-Sumo_(10)_-SIM_(6)_-GFP (Figure 4C; Supplementary movie 1). As expected, multiple RFP-Centrin1 foci are observed on droplets in interphase (Figure 4C,-15 min). As cells enter mitosis, RFP-Centrin1 foci initially remain on the droplet (Fig. 4C, 10 min), but dissociate entirely from it by metaphase (Figure 4C, 30 min). Despite being dissociated from the droplet, ectopic RFP-Centrin1 foci are not eliminated. Instead, they form a cluster inherited by one of the two daughter cells (Figure 4C, 80 min), where they become grouped with resident centrioles during the following G1 (Figure 4C, 90 min). Therefore, ectopic procentrioles dissociate from droplets during mitosis and persist thereafter.

**Figure 4.**
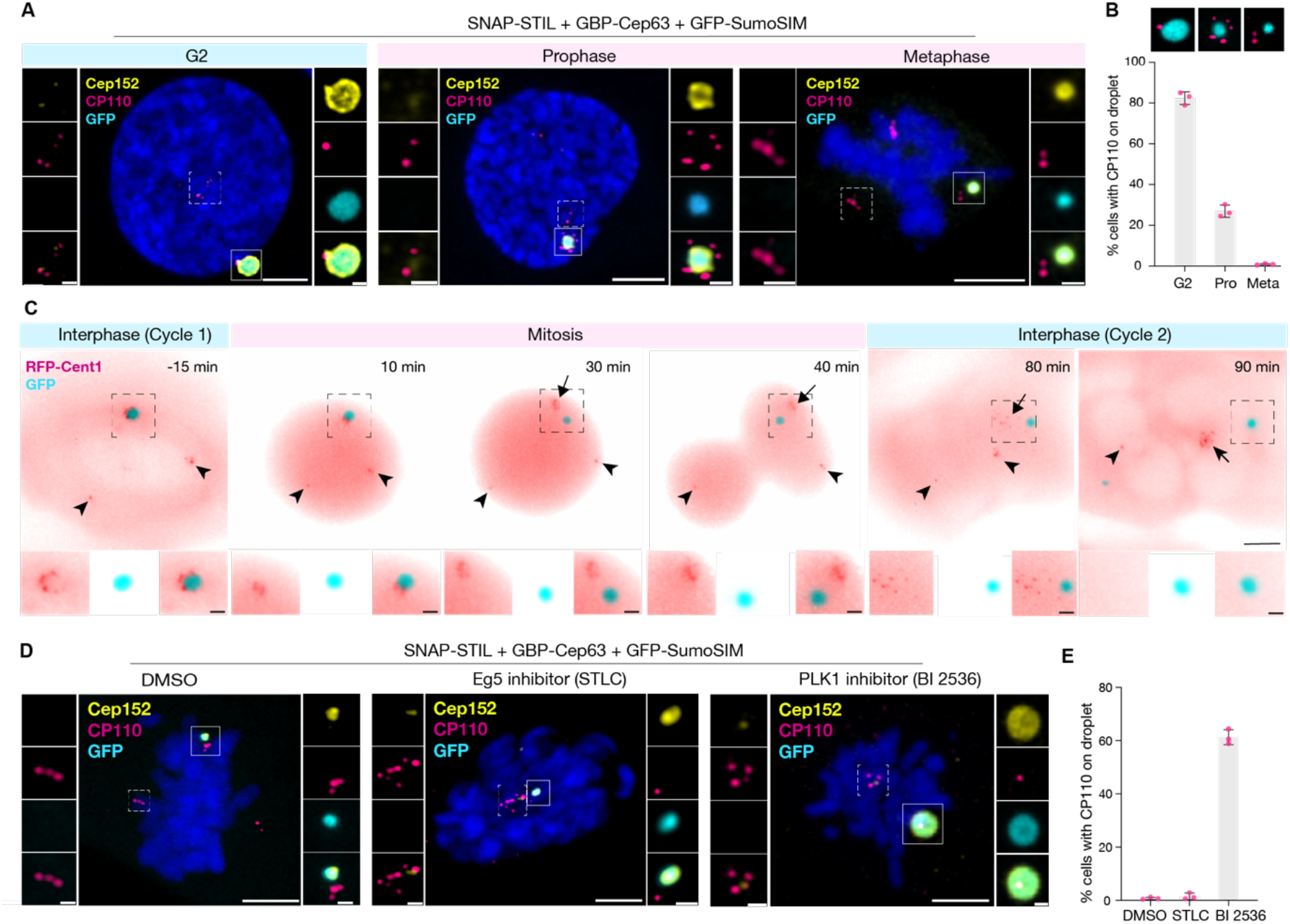
Ectopic procentrioles are released from droplets during mitosis in a PLK1-dependent manner. **A)** Cells expressing SNAP-STIL and GBP-Cep63 in G2, prophase and metaphase, as indicated, stained for Cep152 and CP110. Cell cycle stated was determined by the nuclear signal and position of resident centrioles. Note that CP110 foci remain associated with the droplet until prophase. **B)** Top: magnified images of droplets stained with CP110 as shown in A, in metaphase (left), prophase (middle) and metaphase (right). Bottom: corresponding quantification of percentage of cells where droplets harbor CP110 foci. At least 80 (for G2), 40 (for each prophase and metaphase) cells were scored per experiment in three independent experiments. Bars represent mean +/- s.d. **C)** Live imaging of cells expressing RFP-Centrin1 together with SNAP-STIL and GBP-Cep63 targeted to the droplet marked by GFP. Resident centrioles are indicated with arrowheads. Dashed box highlights the droplet magnified below. Time zero: first instance of cell rounding marking mitosis onset. Note that multiple ectopic procentrioles are released from the droplet beginning in metaphase (30 min). Released ectopic centrioles, indicated by arrow, eventually cluster with the resident centrioles in the following G1. Representative image from 28 cells in 12 independent experiments. **D)** Representative images of cells treated with DMSO, 5 μM STLC or 200 nM BI2536 for 14 hours, and then immunostained for Cep152 and CP110. **E)** Percentage of mitotic cells with persistent CP110 foci on droplets as shown in (D). 50 mitotic cells were scored for each condition per experiment in three experiments.

During the physiological centriole duplication cycle, procentrioles become progressively distanced from resident centrioles, disengaging fully during mitosis through a process that relies notably on the kinase PLK1^29,31,59^. Given the timing of ectopic procentriole dissociation, we addressed whether this process depends on PLK1 activity. To this end, we inhibited PLK1 using BI 2536^60^, finding that this prevents dissociation of ectopic procentrioles from the droplet (Figures 4D, 4E). This is not simply due to a failure of cell cycle progression upon PLK1 inhibition, since cells similarly arrested at prometaphase with the Eg5 inhibitor S-trityl-L-cysteine (STLC)^61^ release ectopic procentrioles from droplets (Figures 4D, 4E). Therefore, mirroring the requirement for procentriole disengagement during the physiological centriole duplication cycle, PLK1 mediates the dissociation of ectopic procentriole from droplets.

### Ectopic procentrioles function as MTOC and seed procentriole formation in turn

We next asked whether dissociated ectopic procentrioles can assemble PCM, nucleate microtubules and generate supernumerary spindle poles. Compatible with this view, we found that ∼57% of cells with a droplet assemble a multipolar spindle (Figure 5A). However, as anticipated from the known impact of excess STIL^15^, overexpressing SNAP-STIL and a Flag tagged version of Cep63 not targeted to the droplet also induces multipolar spindles (Figure 5A). How could spindle poles formed by droplet-dissociated procentrioles be distinguished from those formed merely due to STIL overexpression? We reasoned that upon excess STIL, procentrioles remain next to resident centrioles, such that maturation markers present exclusively on resident centrioles should unequivocally mark such associated supernumerary spindle poles. By contrast, supernumerary spindle poles stemming from ectopic procentrioles dissociated from droplets should not bear such maturation markers. Therefore, we immunostained cells with antibodies against the sub-distal appendage protein ODF2, which marks mature resident centrioles, as well tubulin to mark all spindle poles. As anticipated, we found that control cells overexpressing SNAP-STIL and Flag-Cep63 not targeted to the droplet frequently assemble multipolar spindles, with each pole marked by ODF2 (Figures 5B-C). In contrast, when GBP-Cep63 is directed to the droplet, an ODF2-negative spindle pole often forms in the vicinity of the droplet (Figures 5B, 5C). These ODF2-negative spindle poles frequently harbor a weak focus of γ-tubulin (Figures S6B, S6C), as well as the PCM proteins CDK5RAP2 and Pericentrin (PCNT)^62–64^ (Figure S6D). Together, our data demonstrate that ectopic procentrioles dissociated from droplets act as MTOCs and can organize spindle poles during mitosis.

**Figure 5.**
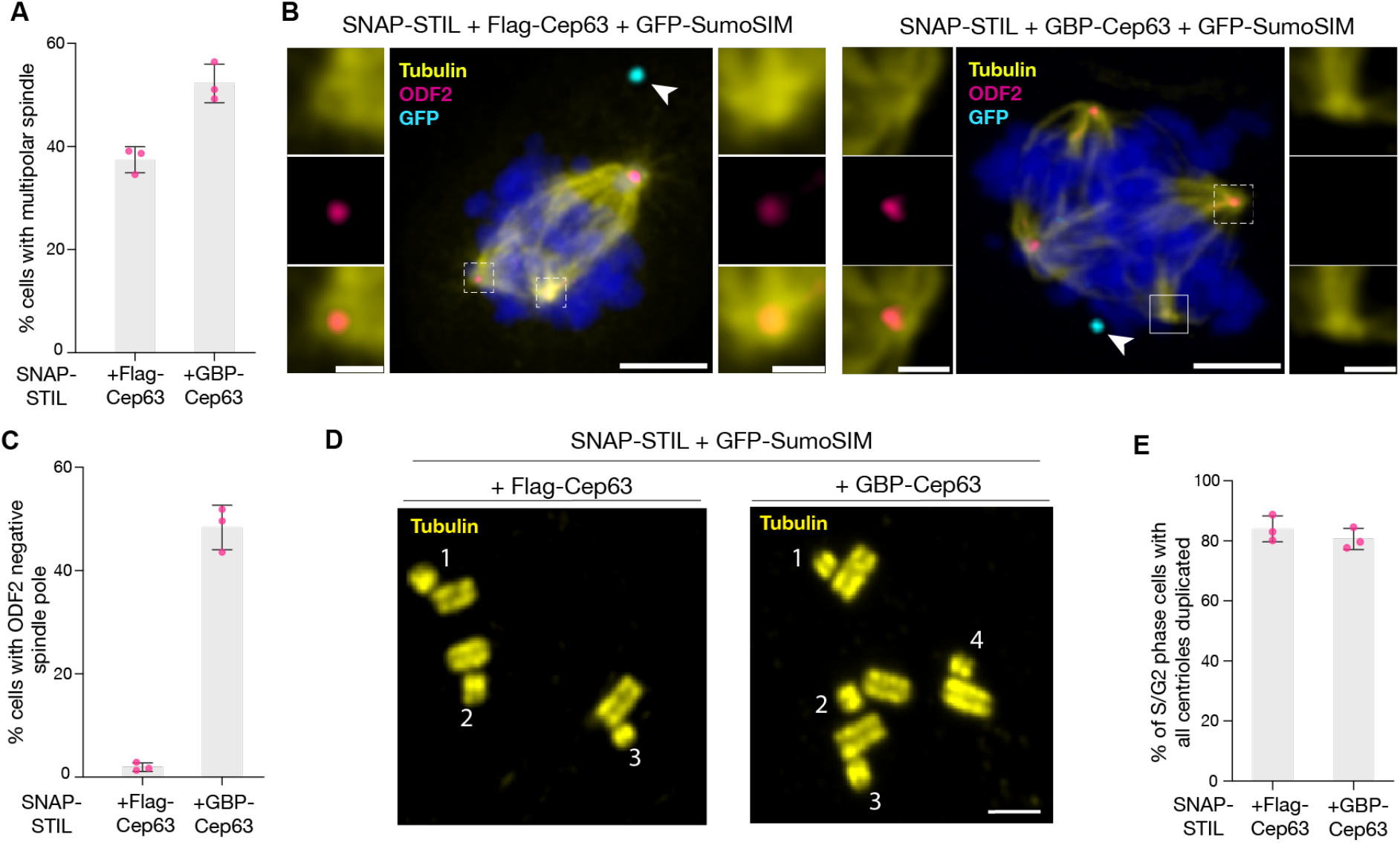
Ectopic procentrioles function as MTOC and seed organelle duplication. **A)** Percentage of mitotic cells with multipolar spindles in cells with SNAP-STIL overexpression, together with either non-targeted (Flag) or droplet targeted (GBP) Cep63. At least 50 cells were scored for each condition per experiment in three experiments. **B)** Representative mitotic cells stained for α/β-tubulin and ODF2 in conditions outlined in (A). Arrowheads point to droplet, dashed boxes to ODF2-positive spindle pole formed around resident centriole, and solid white box to ODF2-negative spindle pole formed by ectopic procentrioles released from the droplet. **C)** Percentage mitotic cells with ODF2 negative spindle poles in conditions outlined in (A). At least 50 cells were scored for each condition per experiment in three experiments. **D)** Representative tubulin U-ExM images of resident centrioles in cells with SNAP-STIL overexpression, together with either non-targeted (Flag) or droplet targeted (GBP) Cep63. Cells were also stained with antibodies against CCDC77 (not shown). Note that all resident centrioles in the cell (3 on the left, 4 on the right, as indicated by numbers) seed one procentriole. Note also that to study centriole fate in the cell cycle following release from the droplet, cells were analyzed 72 hours after STIL and Cep63 induction here, as opposed to 48 hours in the rest of the study. **E)** Corresponding quantification of percentage of S/G2 cells (as indicated by them having at the least one procentriole) in which all resident centrioles have an associated procentriole, as illustrated in (D). Note that in both Flag-Cep63 and GBP-Cep63 conditions, ∼20% of cells have at least one resident centriole without procentriole, likely reflecting asynchrony in organelle biogenesis. 40 cells were counted for each condition in three experiments. Scale bars: 5 μm in low magnification views, 1 μm in insets. Scale bar in D: 250 nm.

We next asked whether procentrioles released from the droplet during mitosis can be converted into full-fledged centrioles that seed procentriole formation in turn in the following cell cycle, as suggested by the presence on ectopic procentrioles of the critical conversion protein Cep295 (see Figure 3H). We reasoned that if this were the case, then all centrioles in the following cell cycle should seed a new procentriole, irrespective of whether they had been released during the preceding mitosis from the droplet or from resident centrioles. To test this prediction, we used U-ExM and tubulin immunostaining to examine all centriolar cylinders present in the following cell cycle, focusing on those cells in which at least one procentriole is present, thereby ensuring that only those cells competent to form procentrioles are analyzed. As shown in Figures 5D and 5E, we found that all resident centrioles are associated with a procentriole in ∼80% such cells overexpressing SNAP-STIL and GBP-Cep63, as in control cells expressing SNAP-STIL and Flag-Cep63. Since cells overexpressing SNAP-STIL and GBP-Cep63 all inherit ectopic procentrioles released from the droplet during the preceding mitosis, it follows that these must have been converted to centrioles able to seed procentriole formation in turn. We also observed that there is a higher percentage of cells with more than two resident centrioles with procentrioles upon overexpression of SNAP-STIL and GBP-Cep63 compared to SNAP-STIL and Flag-Cep63, as anticipated from the former having inherited procentrioles from the droplet in addition to from resident centrioles (Figure S6E).

Overall, these findings indicate that droplet derived procentrioles are functionally equivalent to endogenous procentrioles in their ability to organize spindle poles and seed procentriole assembly in the following cell cycle.

### Surface Cep63 directs *de novo* biogenesis

We sought to deploy the engineered droplet platform to probe centriole biogenesis in an experimental setting in which resident centrioles are lacking, notably to address whether competition for shared resources could explain the observed reliance on excess STIL for full-fledged organelle biogenesis. We reasoned that in the absence of competition from resident centrioles, GBP-Cep63 concentration on the droplet could suffice to trigger procentriole assembly at this ectopic location, without excess STIL. Moreover, we reasoned that the droplet platform should enable us to test whether formation of multiple procentrioles during *de novo* biogenesis reflects dilution of components between several sites. If this were the case, then centriolar components should concentrate and form centrioles *de novo* exclusively at the droplet and not elsewhere in the cell. To conduct these experiments, we treated cells chronically with the reversible PLK4 inhibitor Centrinone, such that the vast majority of cells lacked resident centrioles^65^. Centrinone was then washed out and cells fixed 16 hours thereafter for immunostaining with antibodies against Cep152, CPAP, CP110 and Cep295 (Figures 6A-C). As anticipated, without doxycycline, numerous foci bearing these proteins form *de novo* throughout the cytoplasm (Figure 6B). In contrast, upon doxycycline-mediated induction of GBP-Cep63, we found that Cep152, CPAP, CP110 and Cep295 foci form exclusively on the droplet, indicating the presence of full-fledged procentrioles (Figures 6C, 6D). Moreover, analysis with U-ExM established that SAS-6 and centriolar microtubules are present in these droplet-associated foci (Figure 6E). Importantly, we found also that such droplet derived structures could seed the formation of procentrioles in turn in the following cell-cycle, further demonstrating that droplet-based assemblies mature into full-fledged centrioles (Figure S7A).

**Figure 6.**
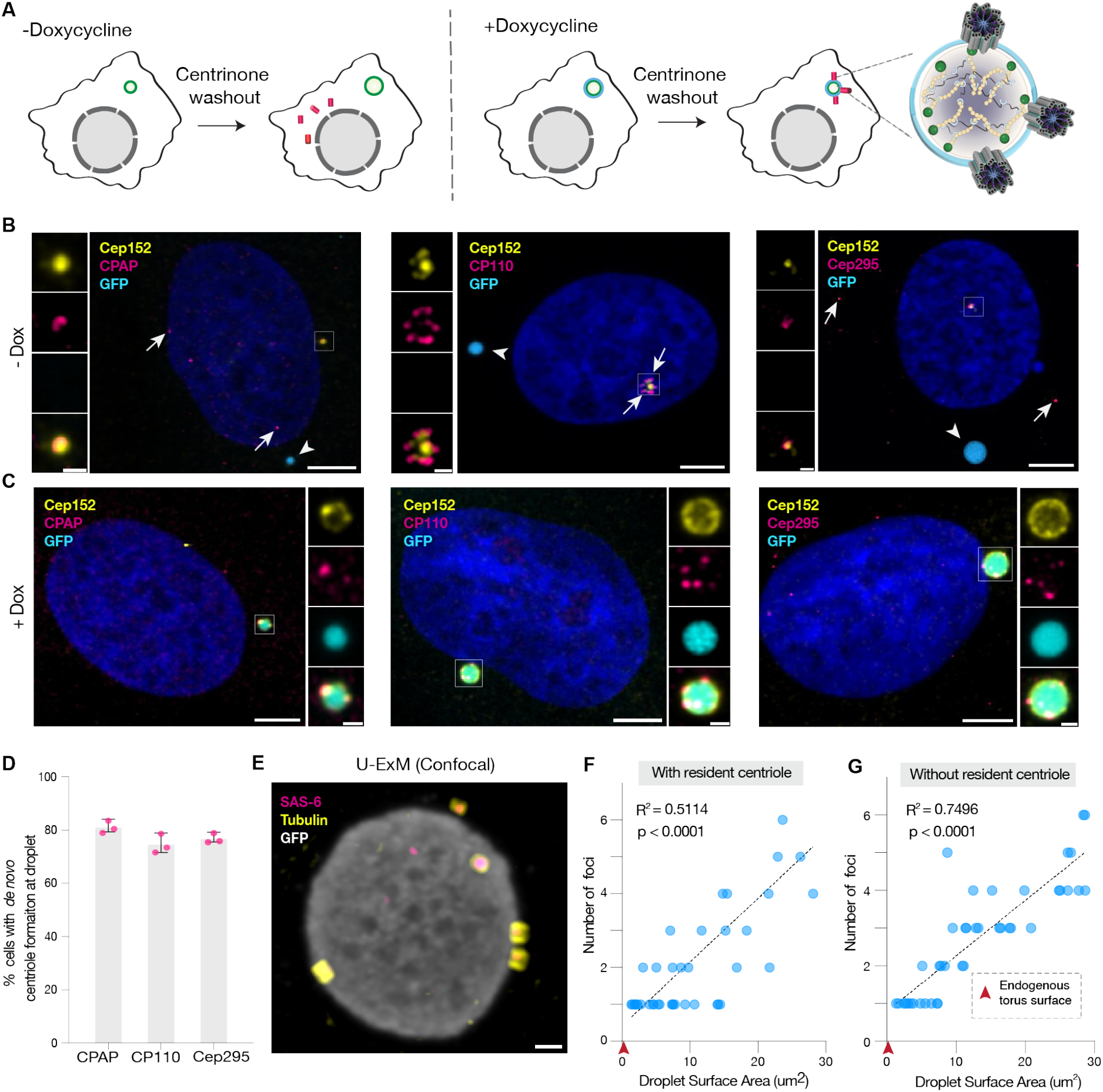
Droplet-targeted Cep63 is sufficient for de novo centriole formation. **A)** Schematic representation of experiment without resident centrioles. Stable cell lines constitutively expressing droplets were treated with Centrinone for seven days, followed by Centrinone washout for 16 hours to induce *de novo* centriole formation without (left) or with (right) doxycycline addition to induce GBP-Cep63 expression. **B, C**) Cells without (B) or with (C) doxycycline induction of GBP-Cep63, immunostained with Cep152 together with indicated centriolar proteins. Arrowheads marks droplet. Note formation of multiple foci throughout cytoplasm in the absence of doxycycline (B, white arrows), versus formation of CPAP, CP110 or Cep295 foci exclusively on the droplet upon Cep63 targeting to the droplet (C). **D)** Percentage of cells where droplets co-localize with indicated centriolar proteins as a percentage of total number of cells undergoing *de novo* centriole formation in the population scored with the same centriolar protein. At least 50 cells were scored per experiment in three experiments. Bars represent mean +/- s.e.m. **E)** Representative max-projected U-ExM image of droplets stained for SAS-6 and α/β-tubulin in *de novo* experimental setting. **F, G**) Number of foci formed on the droplet as a function of droplet surface area with (F) or without (G) resident centrioles. The red arrowhead indicates the surface area of the endogenous torus, as reported before^67^. n=30 cells with a single droplet were analyzed. R^2^ indicates goodness of fit of the linear regression line. P<0.0001 for both F and G and was determined using a t-test for regression co-efficient. Scale bars in B and C: 5 μm in low magnification views, 1 μm in insets. Scale bar in E: 250 nm.

We next capitalized on the size variability of the engineered droplet platform to explore the contribution of torus surface dimensions in regulating procentriole formation. Theoretical considerations have suggested that the single site of procentriole formation at the resident centriole is determined by a Turing-like reaction-diffusion process, the length scale of which affects the number of resulting procentrioles^66^. This implies that the number of procentrioles depends on the size of the surface available to trigger the assembly process. Whereas the invariable diameter of the torus at resident centrioles prevents testing this prediction experimentally, we reasoned that the size variability afforded by the engineered droplets should enable us to do so. As shown in Figure 6F, we found a linear correlation between the surface area of the droplet and the number of ectopic SAS-6 foci when resident centrioles are present (R^2^ = 0.51, p <0.001). Because competition for components at resident centrioles may complicate the interpretation, we likewise asked this question using droplets in Centrinone-treated cells, where such competition is inexistent. Relating the number of CP110 foci to droplet size revealed a stronger linear correlation in this case (Figure 6G; R^2^ = 0.74, p <0.001). Intriguingly, extrapolating from the relationships between droplet surface area and ectopic assemblies, we uncovered that the surface area of the endogenous torus is expected to support formation of a single procentriole (Figures 6F, 6G, red arrowheads)^67^. This relationship pertains only to platform size and not to protein amount present on the surface, as illustrated by mean Cep152 intensities not scaling with CP110 foci number (Figure S7B).

Together, these findings suggest that the dimensions of the torus surface contribute to procentriole number control, illustrating the power of the novel engineered platform to test principles of centriole assembly.

## Discussion

Centrioles are unique amongst cellular organelles in being formed at each cell cycle in a single copy, next to a resident centriole. The mechanisms ensuring the assembly of this evolutionarily conserved mesoscale complex in precisely one copy and at precisely one location constitute an important open question in biology. Here, we engineered a phase-separated droplet platform that enables targeting of individual components to an ectopic cellular location to ask two related questions: first, can a synthetic centriole be reconstituted at an ectopic cellular location; second, if so, which components are sufficient to achieve this feat.

Our engineered platform fulfills two critical prerequisites required to reconstitute low-abundance mesoscale complexes such as centrioles: high valency and low copy number. High valency is essential to target a substantial fraction of the cellular protein pool ectopically, whereas low copy number is needed to avoid diluting between multiple locations proteins present in low amounts. Directing proteins to high valency sites on mitochondria or the plasma membrane has proven useful in other contexts^68,69^, but this results in substantial dilution of low amount proteins. One such protein is STIL, which is present at merely ∼5000-7000 copies per cell^70^. The platform developed here is modular in nature, such that it can be leveraged to ectopically target proteins of other mesoscale macromolecular complexes, including the kinetochore or the nuclear pore. Moreover, by making droplet targeting inducible, for instance with the CatchFire system^71^, this platform can be further engineered to address the kinetics of assembly reactions.

Deploying this platform to investigate centriole biogenesis revealed that targeting the torus protein Cep63 to the droplet is sufficient in and of itself to trigger ectopic assembly of a naked cartwheel. Moreover, Cep63 is sufficient to generate full-fledged procentrioles provided a limiting STIL-CPAP interaction is alleviated or resident centrioles are absent. The importance of STIL echoes findings in the accompanying manuscript with functionalized beads in Drosophila embryos, where homologues of Cep152, STIL, SAS-6, Cep295 and CPAP are each sufficient to seed Centrosome Like Particles (CLiPs) with microtubule organization and self-propagating activities. The interaction between STIL and CPAP also plays a pivotal role in that system, since a STIL/Ana2 mutant unable to bind CPAP/Sas-4 cannot form CLiPs. Intriguingly, Drosophila lacks a Cep63 homologue, raising the possibility that another component, perhaps Cep152/Asl, takes on its role in the fly.

Our findings indicate that Cep63 is unique amongst torus protein in mammalian cells in enabling singly to form a naked cartwheel. Interestingly, during multiciliogenesis, the Cep63 paralogue Deup1 forms spherical structures that guide formation of several procentrioles, in a manner reminiscent of the situation in cycling cells endowed with synthetic droplets^42,72,73^. Given the importance of Cep63 in ensuring procentriole formation in the proper location and with the proper number, cytosolic levels of this protein ought to be regulated to prevent illegitimate procentriole assembly, which may help explain why autophagosomes specifically eliminate cytosolic Cep63 foci in mouse embryonic fibroblasts^74^.

The mechanisms instructing procentrioles to form in the proper location and number have intrigued scientists since the centriole duplication cycle was revealed by EM decades ago^75,76^. Nucleic acids have been postulated to be present at centrioles and to convey such instruction, although this proposal lacks experimental backing^77^. Alternatively, it has been suggested that the resident centriole could act as a template for cartwheel formation, thereby explaining the seeding of one procentriole in its vicinity^78^. A simpler possibility is that the torus surface is critical for instructing the assembly site and determining procentriole number. By distancing the site of procentriole formation from resident centrioles, our work provides a test of this possibility. We show that procentriole number scales linearly with torus surface area and, extrapolating from this relationship, that the surface area of the endogenous torus is optimal for supporting assembly of a single procentriole. Intriguingly, the dimensions of the torus, and hence its surface area, are set by resident centriole size, with the innermost torus components Cep57 binding to the centriolar microtubule wall^3,79^. We propose that this mechanism constitutes a closed-loop in which the dimensions of the torus ensure singularity of procentriole formation, with the size of the resulting resident centriole in turn dictating torus dimensions, thereby ensuring organelle number homeostasis across cell generations.

### Limitations of the study

Our study demonstrates that concentrating Cep63 is sufficient to trigger procentriole assembly. However, we have not determined which part of the Cep63 protein is sufficient to trigger the assembly reaction. In addition, our findings with Cep63 do not exclude that other components may do so as well, which can be tested in the future using our versatile engineered platform.

We also did not investigate the kinetics of PLK4, STIL and SAS-6 focusing on the droplet surface, nor the underlying mechanism of such focusing. Future work with live-cell imaging and inducible targeting to the droplet can serve to clarify this point.

Although we have established that droplet derived procentrioles are functional in organizing PCM and spindle poles during mitosis, as well as in seeding procentriole formation at the following cell cycle, we have not tested whether they can act as basal bodies during primary cilium formation, as we have used U-2 OS cells, which do not have this capacity. Future investigations in cell lines that can form primary cilia will be informative in this regard.

## Acknowledgements

We thank Friso Douma, Anoek Friskes, Juan Manuel Garcia Arcos and Gabriela Garcia-Rodriguez for constructive comments on the manuscript, as well as all Gönczy lab members for discussions. We also thank Emmanuel Derivery, Mark Van Breugel, Ruud Hovius and Gislene Pereira for kindly sharing reagents, as well as George Hatzopoulos, Graham Knott, Francisco Palumbo, Gabriele de Simone, Cédric Pourroy and Camilla Brückmann de Mattos for expertise.

## Author contributions

Design of the study: S.A. and P.G. Data acquisition: S.A. Data analysis: S.A. Manuscript preparation: S.A. and P.G. Supervision: P.G. Funding acquisition: S.A. and P.G.

## Methods

### DNA constructs

The original RFP-Sumo_(10x)_-SIM_(6x)_ construct^48^ was obtained from Addgene (cat. 122026) and modified with restriction enzymes Age1 and BspE1 to replace RFP with GFP. The version of GFP used in this work is eGFP and is referred simply as ‘GFP’ throughout. ORFs for all centriolar torus proteins were originally sourced from Lukinavičius et al.^8^ or the CSBB-Broad Institute Lentivirus expression library (for Cep57L1). Each ORF was subcloned to be flanked by Fse1 and Asc1 sites for convenient shuttling.

All ORFs were cloned into a pCDNA/FRT vector modified to include the MXS chaining system^80^. This allows for polycistronic expression of multiple ORFs, each with its own promoter and terminator. GFP-Sumo_(10x)_-SIM_(6x)_ was expressed constitutively with a CMV promoter, whereas all centriolar protein ORFs were placed under the control of doxycycline-inducible third generation Tet promoter. All vectors were verified by whole plasmid sequencing.

### Cell culture

Control U-2 OS (Sigma, cat. 2022711) and U-2 OS FlpIn cells^81^ were cultured in DMEM (Thermo Fisher Scientific, cat.0565018) with 10% FBS and 100 μg/ml Zeocin for FlpIn maintenance. Cells were cultured in a humidified 5% CO2 incubator at 37°C and split every 3–4 days at 80% confluency. Cells were periodically screened for mycoplasma contamination. For generation of stable cell lines, U-2 OS FlpIn cells were transfected with Lipofectamine 3000 (Invitrogen), with a 1:3 ratio of pOG44 and pFRT plasmid encoding the construct of choice, according to the manufacturer’s instructions. Cells with pFRT integration were selected for two weeks with 500 μg/ml hygromycin and cells with high expression of the construct selected by FACS using GFP (GFP-SumoSIM is expressed as part of the same polycistronic vector as mentioned above). Cells were maintained thereafter in 100 μg/ml hygromycin. Cell lines used in this study are listed in Supplemental Table 1.

For siRNA experiments, both negative control (silencer select, Invitrogen, 390843) and siRBM14 (silencer select, Invitrogen, s20406) were transfected using RNAiMax (Invitrogen) at 20 pmol for 72 hours, according to the manufacturer’s instructions. 1 μg/ml doxycycline was added 24 hours after siRNA transfection.

For all assays, cells were induced with 1 μg/ml doxycycline for 48 hours before fixation, except for reduplication assay (Figures 5D, 5E) where cells were fixed 72 hours thereafter. For PLK1 inhibition experiments (Figure 4D), 200 nM BI 2536 (MCE, cat. HY-50698) was added for the last 14 hours before fixation. As control, cells were treated with either 5 μM STLC (Sigma, cat 164739) or 0.001 % DMSO for the same duration.

For *de novo* biogenesis experiments (Figure 6), cells were treated with 125 nM Centrinone B (MCE, cat. HY-18683) for 7 days before seeding in 6-well plates and doxycycline addition. Centrinone was removed by extensive washing with PBS after 32 hours and cells fixed 16 hours later (corresponding to 48 hours after doxycycline addition).

### Fixation and immunofluorescence

Cells were grown on 12 mm diameter untreated glass coverslips and fixed with 4% paraformaldehyde (PFA) in PBS for 10 min at room temperature. Excess PFA was quenched with fresh NaBH4 (2 mg/mL in PBS, SigmaAldrich, cat. 213462) for 5 min and post-fixed with 100% methanol at-20°C for 7 min before rehydration and storage in PBS. This fixation method allowed for preservation of droplet structure while allowing for staining of most centriolar proteins and was followed for all experiments except for Figure S4 where cells were directly fixed with 100% methanol.

For immunostaining, cells were permeabilized with 0.5% (v/v) Triton-X100 in PBS for 15 min, washed in PBS and 0.05% (v/v) Tween 20 (PBST), before blocking for 30 min in PBST supplemented with 3% BSA. All antibodies were diluted in the blocking buffer and incubated overnight at 4°C for primary antibodies and 45 min at room temperature for secondary antibodies. Primary antibodies are listed in Supplemental Table 2, secondary antibodies in Supplemental Table 3. After secondary antibody labelling, cells were stained with 1 μg/ml Hoechst 33258 (Sigma) in PBS for 5 min to mark DNA, prior to mounting in Fluoromount-G (ThermoFisher Scientific, cat. 00-4958-02).

### Live-cell imaging

U-2 OS cells stably expressing RFP-Centrin1 were plated on glass bottom µ-Dish 35 mm Quad dishes (Ibidi, cat. 80416) and transfected with a single polycistronic vector constitutively expressing GFP-SumoSIM and doxycycline inducible GBP-Cep63 and SNAP-STIL. 24 hours after induction with 1μg/ml doxycycline, the medium was replaced with live-cell imaging medium containing Fluorobrite DMEM (Thermo Fisher Scientific, cat. A1896701), GlutaMAX (100x, 1/100, Thermo Fisher Scientific, cat. 35050061), 10% FBS and Penicillin-Streptomycin (Merck, 100 U/mL, cat. P0781).

Images were acquired every 10 min for at least 24 hours using a Zeiss Observer D1 wide-field microscope with a 63x oil immersion objective (NA 1.40) and equipped with an Andor Zyla 4.2p EMCCD camera. Z-sections were taken every 400 nm for a total of 8um Z volume. Cells were imaged at 37°C with 5% CO2. Both the stage-top chamber and objective heating were controlled by Okolab instruments.

### Expansion

Cells were expanded according to Gambrotto et al.^82^. Briefly, coverslips fixed as above were crosslinked in a 6-well plate in Acrylamide/Formaldehyde solution (1% Acrylamide and 1% Formaldehyde in PBS) overnight at 37°C. For gelation, coverslips were incubated in 40 μl monomer solution containing 19% (wt/wt) Sodium Acrylate, 10% (wt/wt) Acrylamide, 0.05% (wt/wt) BIS in PBS supplemented with fresh 0.5% Tetramethylethylenediamine (TEMED) and fresh 0.5% Amonium Persulfate (APS) on a pre-cooled piece of parafilm for 1 hour at 37°C in a moist chamber in the dark. To release the gels from the coverslips, coverslips were incubated for 15 min in denaturation buffer (200 mM SDS, 200 mM NaCl and 50 mM Tris pH = 9) followed by incubation for 1 hour on a 95°C hot plate with periodic agitation in fresh denaturation buffer. Post-denaturation gels were expanded by incubation in water twice for 15 min each, followed by overnight incubation at room temperature. The expansion factor was quantified by carefully measuring the diameter of the expanded gel with a caliper and dividing this by the initial size (12 mm). After expansion, gels were cut in pieces with a home-made 3D printed punching device. Gels were either stored in water at 4°C or shrunk by incubating with PBST for 15 min. Staining was performed overnight with primary antibodies diluted in blocking buffer at room temperature. Gels were further washed 3 times in PBST for 10 min each, before incubation with secondary antibodies also diluted in blocking at 37°C in the dark for 2 hours. Finally, to label nuclei, gels were stained with 1 μg/ml Hoechst 33258 (Sigma) in PBS for 5 min. For re-expansion, gels were washed thrice for 15 min each in distilled water. For imaging, gels were placed on poly-D-lysine coated coverslips, mounted with i-spacers (IS331, SunJin Labs) on a glass slide with minimal water retention.

### Microscopy

All images were acquired on a Leica SP8 upright confocal microscope equipped with a 63X objective (NA 1.4) and hybrid detectors. All immunofluorescence images were acquired with 77nm x-y pixel size and 0.35 Z-steps. For expansion microscopy, images were acquired with 43nm x-y pixel size to enable deconvolution. For quantification of signal intensities, data sets were acquired using a Nikon Crest Spinning disk microscope equipped with a 60X objective (NA 1.42), a Lumencor Celesta 7 NII-AA laser source and a teledyne Kinetix sCMOS camera. For STED imaging (Figure 3K), the Abberior Mirava polyscope was used with the Lightbox software. A 775nm STED laser was used at 60% for 3D depletion with automated gating with 65nm Z step sizes. Images were further deconvolved by intensity using maximum likelihood estimation and reconstructed in the volume viewer of the Lightbox software.

### Image analysis

All images were quantified using raw intensity values after max projection. For Fig 1E, intensities at the droplet were quantified using a custom written automated script which used the GFP channel to locate and threshold (Otsu) around the droplet. This threshold was used as a mask to segment the droplet across all channels and quantify mean intensity and sizes. For the resident centriole, the PCM marker in the far-red channel was similarly used to threshold the centriolar signal. The partition coefficient was calculated as:

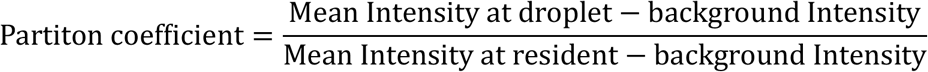

For Figure 2C, to calculate partition coefficient, mean intensities were calculated manually with a 5-pixel wide region of interest, surrounding each protein signal.

For Figures 6F and 6G, the number of foci were counted manually for each droplet. The percentage of cells where a given centriolar protein foci are present on the droplet was calculated by scoring those cells where the centriolar protein in question is also present at resident centrioles (except for Figure 3C, where all cells were scored), which was set to “100%” in these quantifications, allowing us to focus analysis on cells in the correct cell cycle phase for a given centriolar marker to be present.

For quantification of Cep63 or Cep152 diameter, a line profile of 9-pixel width was drawn across the image and distances calculated from peaks found using the “Find Maxima” plugin. For quantification of expansion microscopy data, all images were calibrated with the measured expansion factor and max projected. Length and width were calculated by measuring the full-width half maxima across a line profile fitted with a single gaussian curve for length, and sum of two Gaussian curves for width.

All images were max-projected, and brightness adjusted on the whole image unless otherwise indicated. For representation purposes, immunofluorescence images were smoothened with Gaussian filter with 1-pixel radius. For expansion microscopy data, a similar operation was performed with 2-pixel radius.

### FACS analysis

For nuclear staining, cells were fixed with 70% ethanol at 4°C overnight and stained with 10 ug/ml Propidium Iodide (Sigma-Aldrich, cat. P4170) at 37°C for 2 hours. Cells were filtered through a 35 μm nylon filter and samples analysed on a LSR Fortessa (BD biosciences) analyser with appropriate gating to exclude doublets and dead cells. At least 10,000 cells were analysed for each condition. Cell cycle populations were analysed by FlowJo (version10) software.

## Statistical analysis and reproducibility

Unless otherwise mentioned all experiments were repeated at least three times. No tests were done *a priori* to determine sample sizes. Statistical tests, where appropriate, were performed in Graphpad Prism as mentioned in the respective figure legends. No data was excluded from analysis, and the experiments were not randomized. For Figures 6F and 6G, p value indicates that the slope of the regression line is significantly different than zero, as indicated.

---

**Supplementary Table 1:**

| Reagent | Source |
| --- | --- |
| U-2 OS FlpIn CMV :: GFP-Sumo-SIM | This study |
| U-2 OS FlpIn CMV :: GFP-Sumo-SIM + Tet :: Cep57-GBP | This study |
| U-2 OS FlpIn CMV :: GFP-Sumo-SIM + Tet :: Cep57L1-GBP | This study |
| U-2 OS FlpIn CMV :: GFP-Sumo-SIM + Tet :: Cep57-GBP + Tet :: Cep57L1-GBP |  |
| U-2 OS FlpIn CMV :: GFP-Sumo-SIM + Tet :: GBP-Cep63 | This study |
| U-2 OS FlpIn CMV :: GFP-Sumo-SIM + Tet :: Cep152-GBP | This study |
| U-2 OS FlpIn CMV:: GFP-Sumo-SIM + Tet :: GBP-Cep63 + Tet::SNAP-STIL | This study |
| U-2 OS FlpIn CMV:: GFP-Sumo-SIM + Tet :: Flag-Cep63 + Tet::SNAP-STIL |  |
| U2-OS Tet :: tagRFP-Centrin1 | Keller et al. |
| pTwist FlpIN CMV :: GFP-Sumo-SIM + Tet :: GBP-Cep63 + Tet ::SNAP-STIL | This study |

**Supplementary Table 2:**
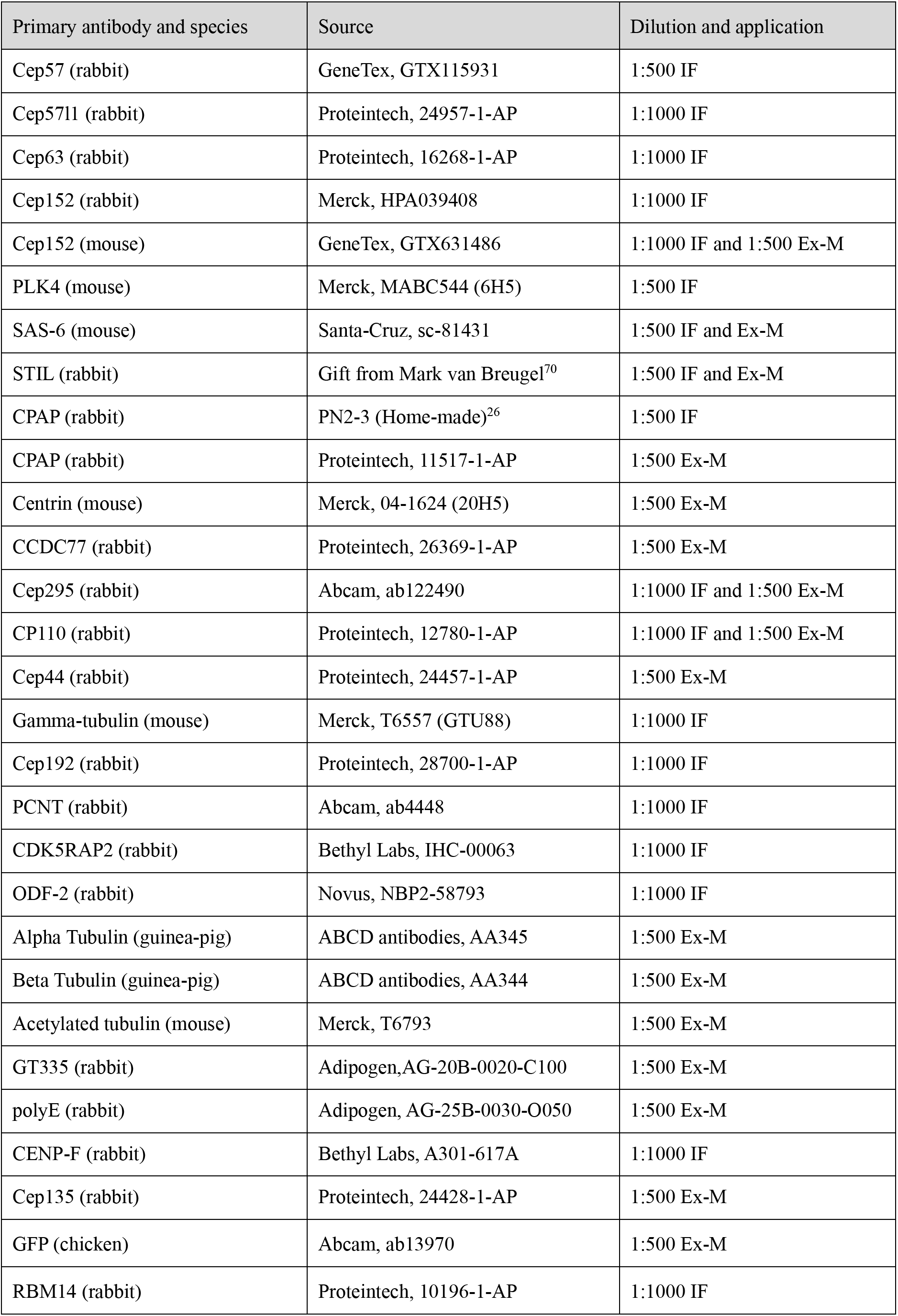

**Supplementary Table 3:**

| Secondary antibody and species | Source | Dilution and application |
| --- | --- | --- |
| Donkey anti-mouse 568 | Invitrogen, A10037 | 1:1000 IF |
| Donkey anti-rabbit 568 | Invitrogen, A10042 | 1:1000 IF |
| Donkey anti-mouse 647Plus | Invitrogen, A32787 | 1:1000 IF |
| Donkey anti-rabbit 647Plus | Invitrogen, A32795 | 1:1000 IF |
| Goat anti-guineapig 488 | Invitrogen, A11073 | 1:500 Ex-M |
| Goat anti-chicken 568 | Invitrogen, A11041 | 1:500 Ex-M |
| Goat anti-mouse 594 | Invitrogen, A-11032 | 1:500 Ex-M |
| Goat anti-rabbit Star 635P | Abberior, ST635P-1001 | 1:500 Ex-M |

**Figure S1.**
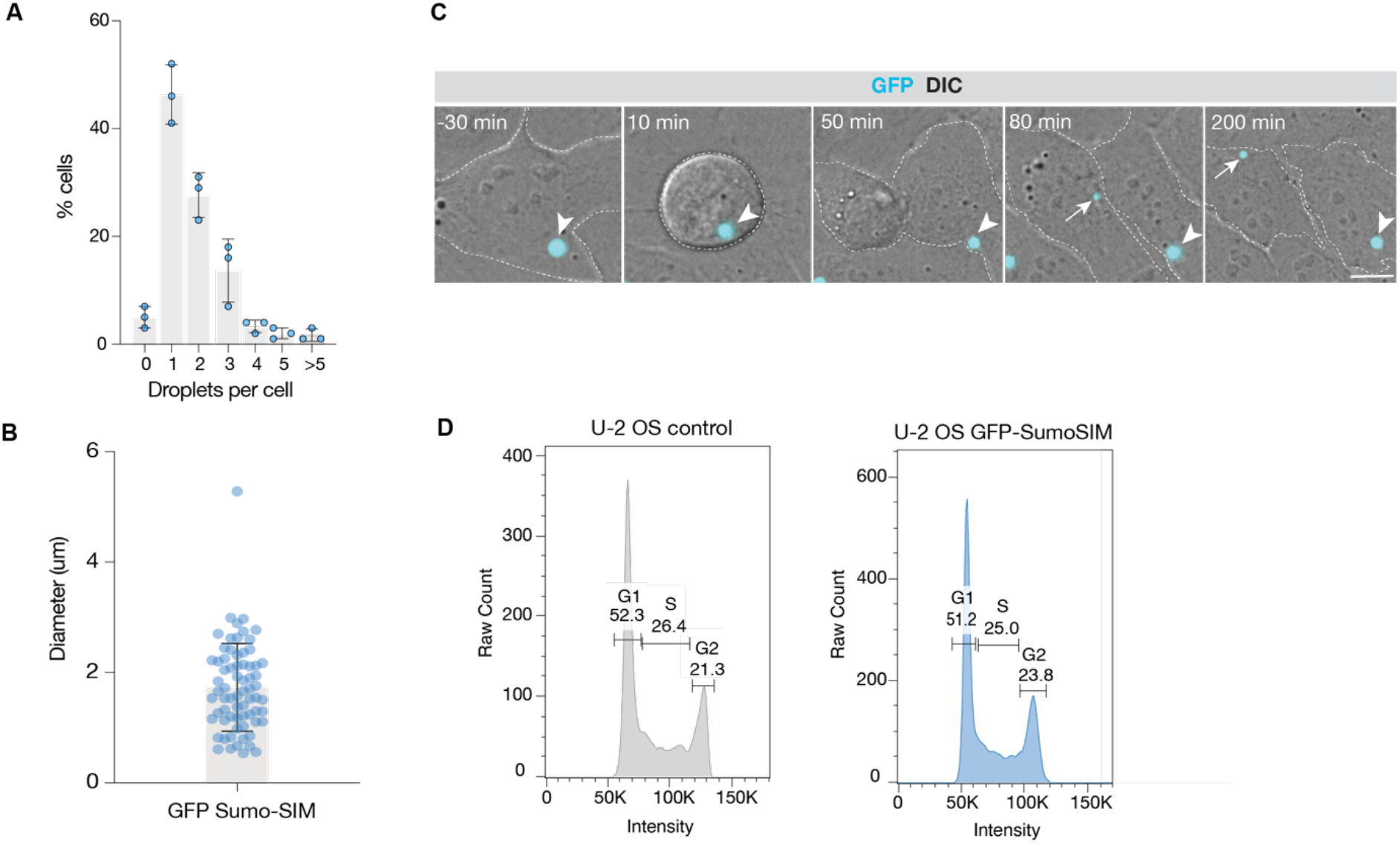
Characterization of cells expressing GFP-Sumo(10x)-SIM(6x) **A)** Percentage of cells with indicated number of droplets. Bars represent mean +/-s.d. of three experiments **B)** Droplet diameter. Each point represents an individual droplet. Bars represent mean +/-s.d. **C)** Montage from live imaging of U-2 OS cell containing a droplet (cyan, arrowhead) undergoing mitosis. Soon after cell division, a new droplet forms in the daughter cell without the inherited droplet, indicated by arrow. Dashed lines represent cell outlines. **D)** FACS profile of DNA distribution of control and droplet expressing cells. At least 10,000 cells were scored per experiment, which was conducted twice, with similar outcomes. Scale bar in B: 5 μm.

**Figure S2.**
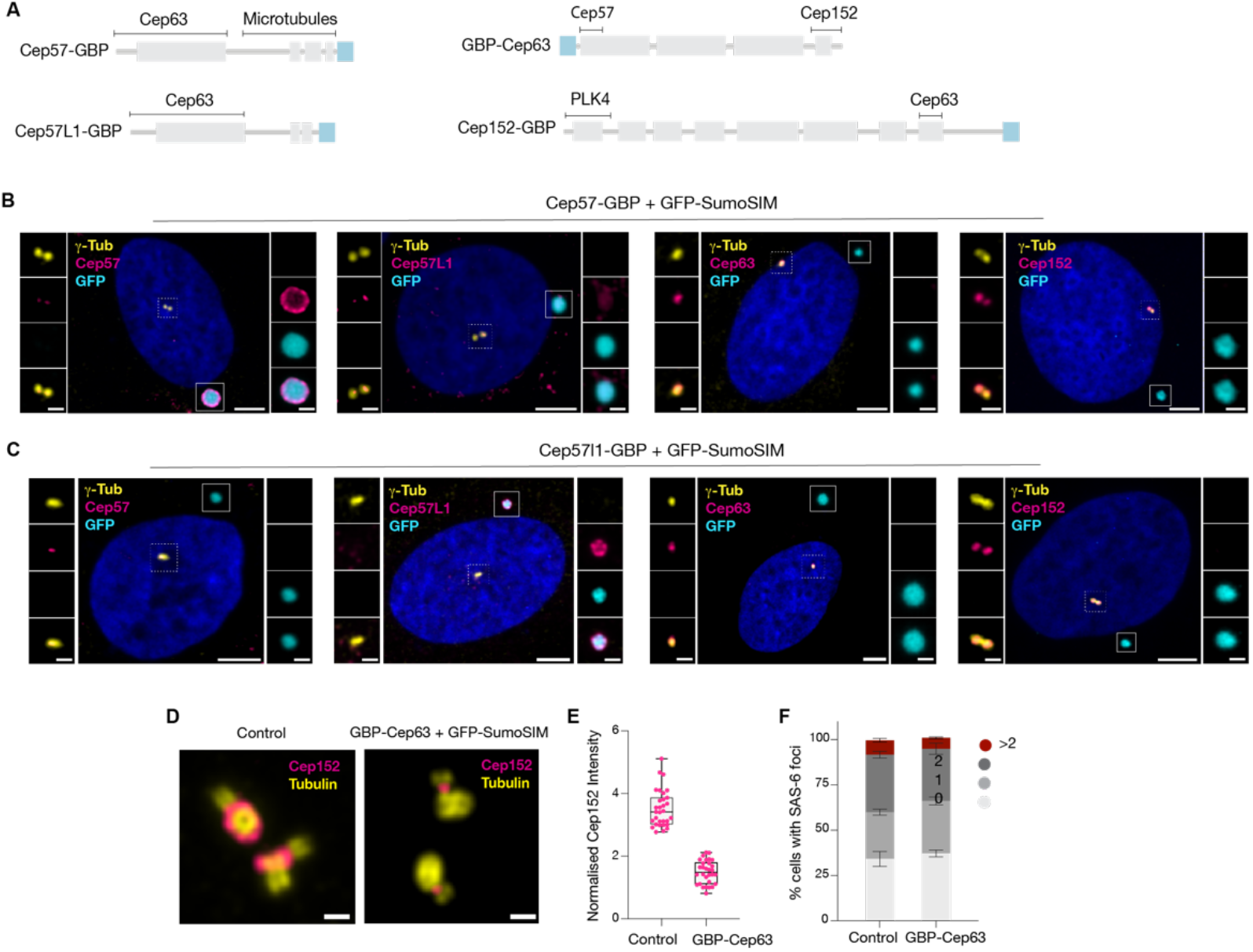
Interdependences of torus proteins following droplet recruitment of individual components. **A)** Torus proteins and their known interaction with other components relevant to the study. Blue segments represent GBP. **B, C**) Representative immunofluorescence images of cells with droplet targeted with indicated torus proteins. γ-tubulin marks resident centrioles. **D)** Representative max-projected U-ExM image of resident centrioles stained for Cep152 and α/β-tubulin in control cells (without doxycycline) or in cells with doxycycline-mediated GBP-Cep63 targeting to the droplet. Note that upon Cep63 droplet targeting, Cep152 is depleted at resident centrioles, without however affecting endogenous procentriole assembly. **E)** Corresponding quantification of Cep152 enrichment at resident centrioles. n=35 cells from two independent experiments. **F)** Quantification of SAS-6 foci number at resident centrioles in either condition outlined in D. 100 cells were counted for both conditions per experiment in three experiments. Scale bars in B and C: 5 μm in low magnification views, 1 μm in insets. Scale bar in D: 250 nm.

**Figure S3.**
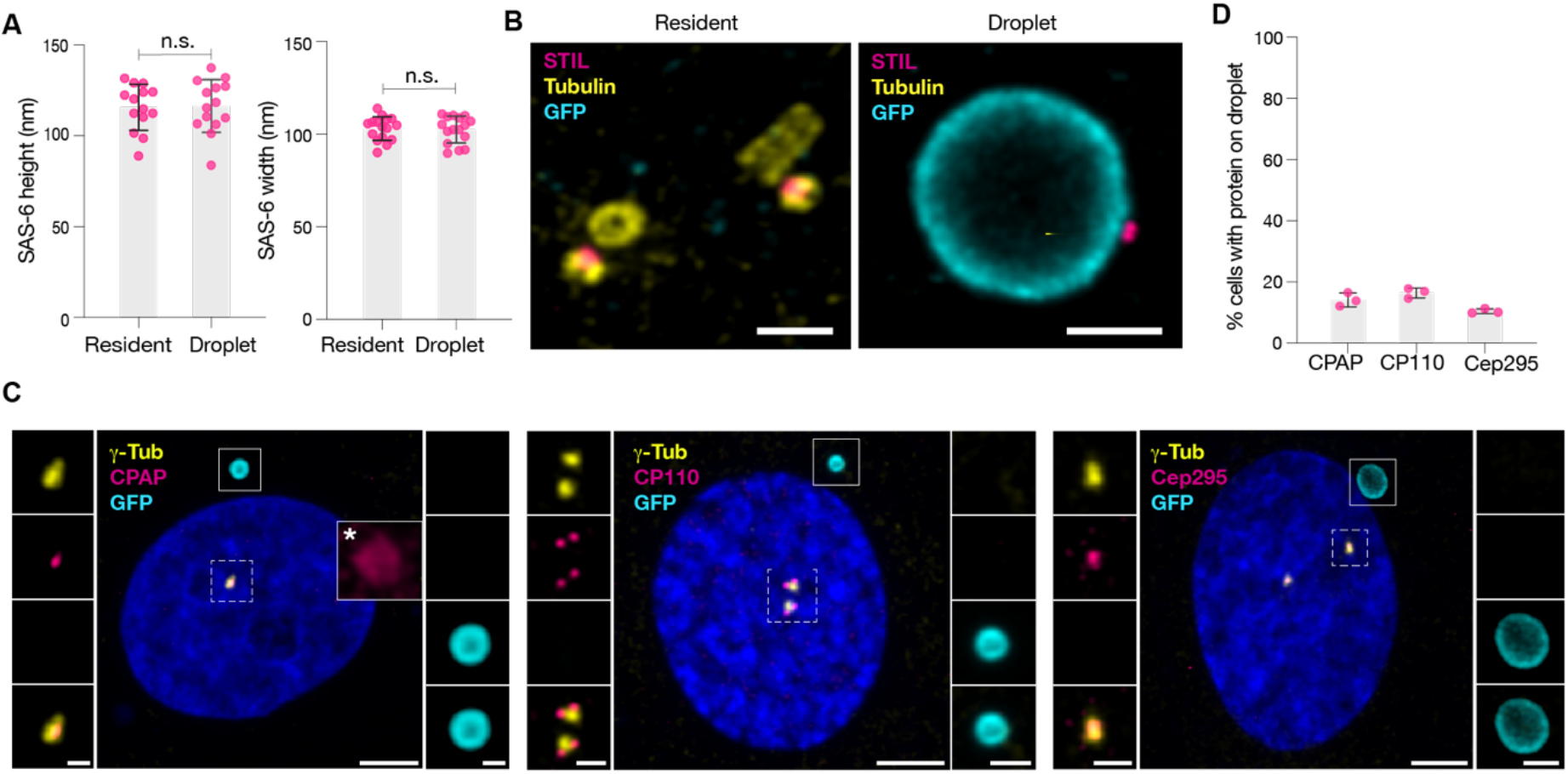
Targeting Cep63 to the droplet results in cartwheel formation but not further procentriole maturation. **A)** Height and width of SAS-6 signal on droplet measured from U-ExM images as shown in Fig. 2F. n=14 (for height) and 16 (for width) from two independent experiments. Each point refers to a cell. Bars represent mean +/-s.d. Significance was tested by paired two-tailed t-test. n.s. indicates p >0.05. **B)** Max-projected U-ExM image of resident centriole and droplet from the same cell stained for tubulin, STIL and GFP. **C)** Representative images of cells with Cep63 targeted droplet stained for the centriolar microtubule binding proteins CPAP, CP110 and Cep295. For CPAP, a higher exposure image of the same droplet (marked by asterisk) is shown to illustrate weak uniform droplet localization and failure to form a focus. **D)** Percentage of cells in which the droplet harbors indicated centriolar proteins on its surface, expressed as a fraction of number of cells where the same centriolar protein marks procentrioles next to resident centrioles. At least 100 cells were counted for each condition per experiment in three experiments. Note that the percentage of cells where markers are present at the droplet is extremely low and may reflect the small fraction of U-2 OS cells known to contain supernumerary centriole in an asynchronous cycling population^65^. Scale bars in B: 250 nm. Scale bars in C: 5 μm in low magnification views, 1 μm in insets.

**Figure S4.**
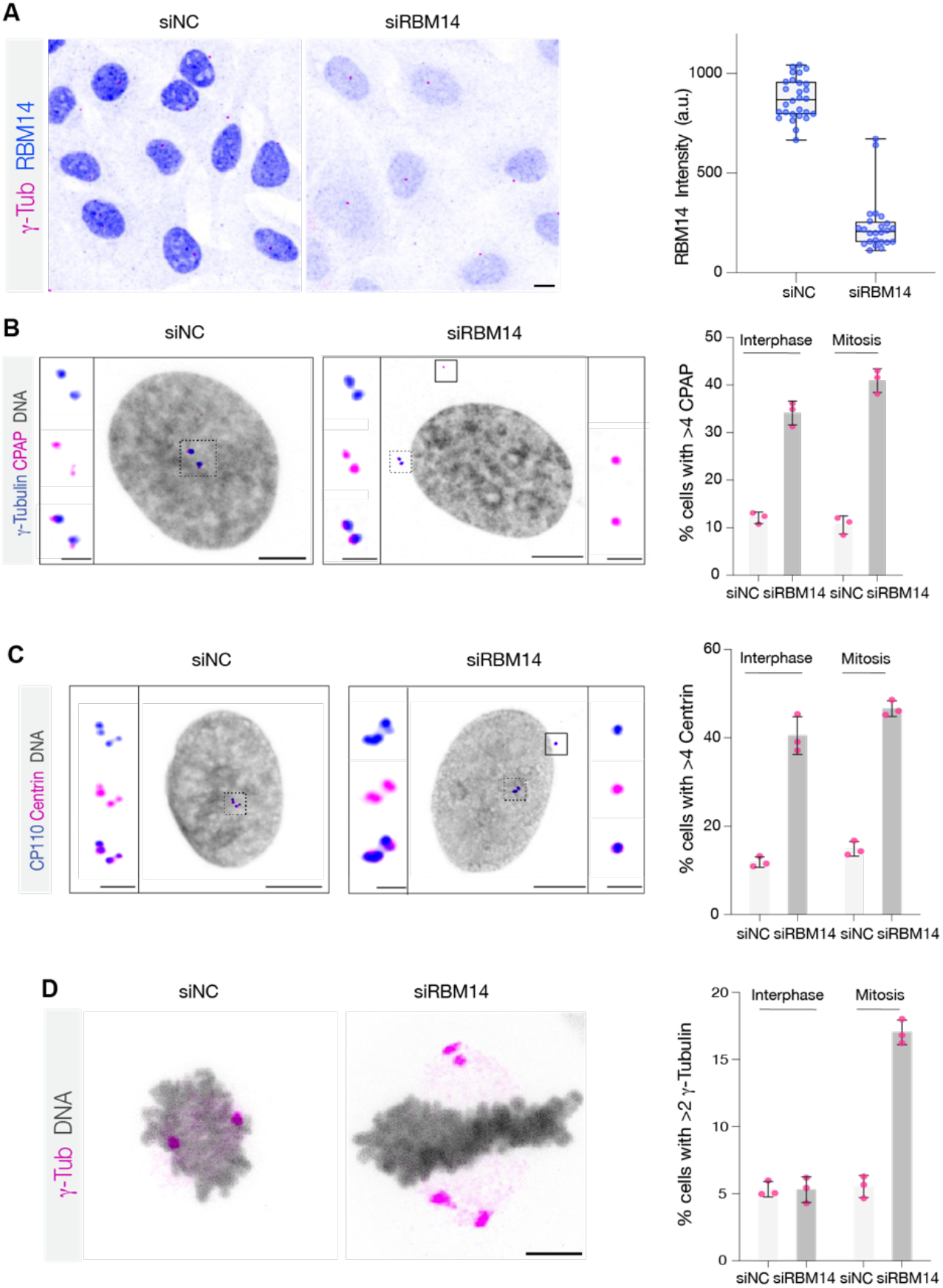
RBM14 depletion causes ectopic centriolar protein assemblies and multipolar mitosis. **A)** Left: representative methanol fixed cells treated with either control siRNA (siNC) or siRNA against RBM14, and then immunostained for RBM14 and for γ-tubulin to mark centrosomes. Right: corresonding quantification of mean RBM14 intensity; each data point represents an individual cell. n=60 cells from three independent experiments. Bars represent mean +/-s.d. **B, C**) Left: representative interphase cells stained for indicated protein, with corresponding quantification on the right. At least 50 cells were scored for each condition per experiment in three experiments. Bars represent mean +/-s.d. **D**) Left: mitotic cells in either control or RBM14 depleted condition, as indicated, and immunostained for γ-tubulin. Note extra γ-tubulin foci upon RBM14 depletion. Right: corresponding quantifications. At least 50 cells were scored for each condition per experiment in three experiments. Bars represent mean +/-s.d. Scale bars: 5 μm in low magnification views, 1 μm in insets.

**Figure S5.**
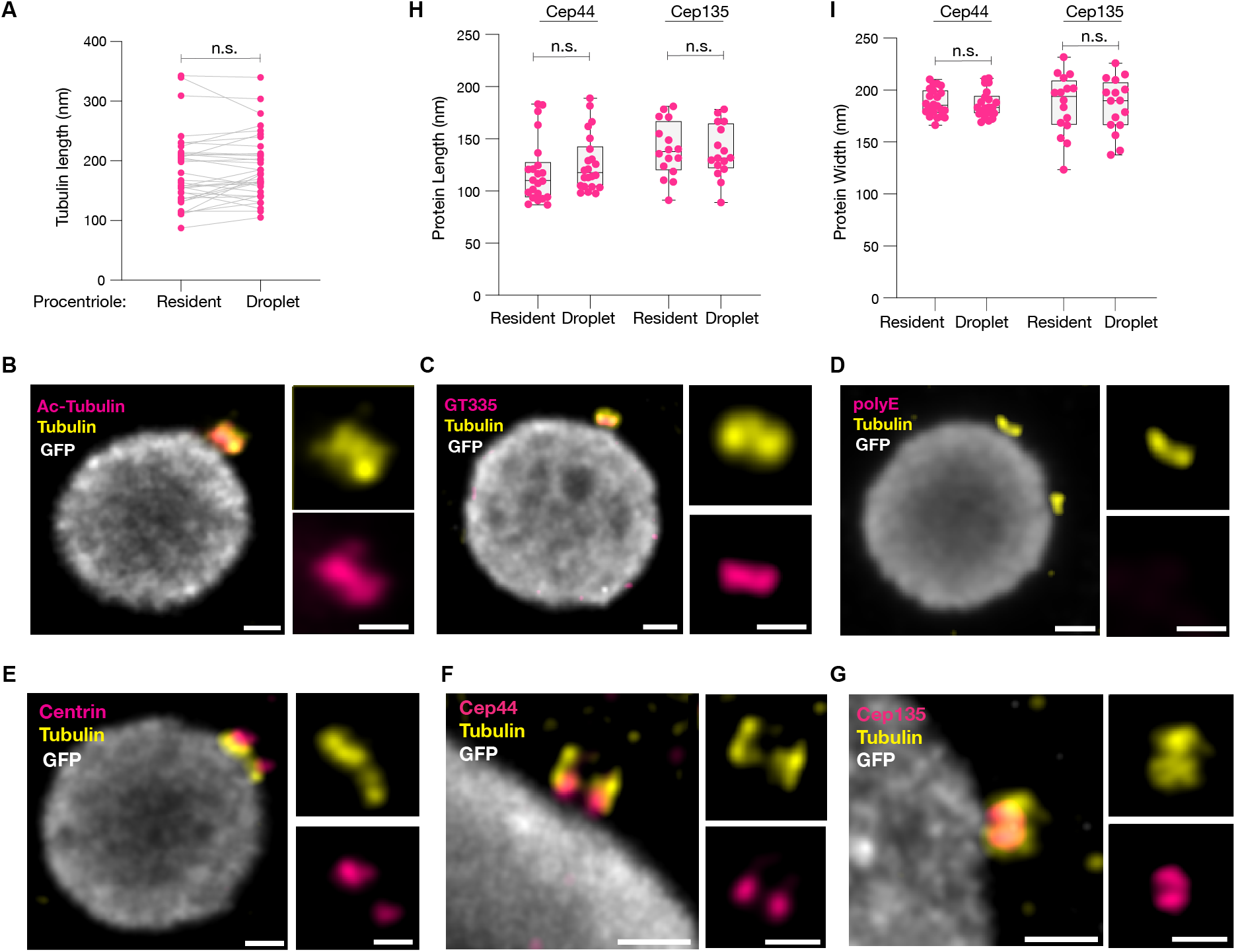
Droplet-assisted ectopic procentrioles have features characteristic of mature procentrioles. **A)** Tubulin length shown in Fig. 3E replotted as a “before-after” graph for paired comparisons. Statistical significance was tested by two-tailed paired t-test. p =0.2128. **B-G**) Representative max-projected U-ExM images of droplets stained for tubulin and indicated microtubule modifications (B-D) or centriolar proteins (E-G). **H, I**) Length and width of Cep44 and Cep135 signals at the resident centriole compared to the droplet. Each data point represents an individual procentriole, boxes represent interquartile ranges, horizontal line indicates median and whiskers represent minima and maxima. n=22 (Cep44) and 16 (Cep135) pooled from two independent experiments. n.s. indicates p >0.05, measured using paired two-tailed t-test.

**Figure S6.**
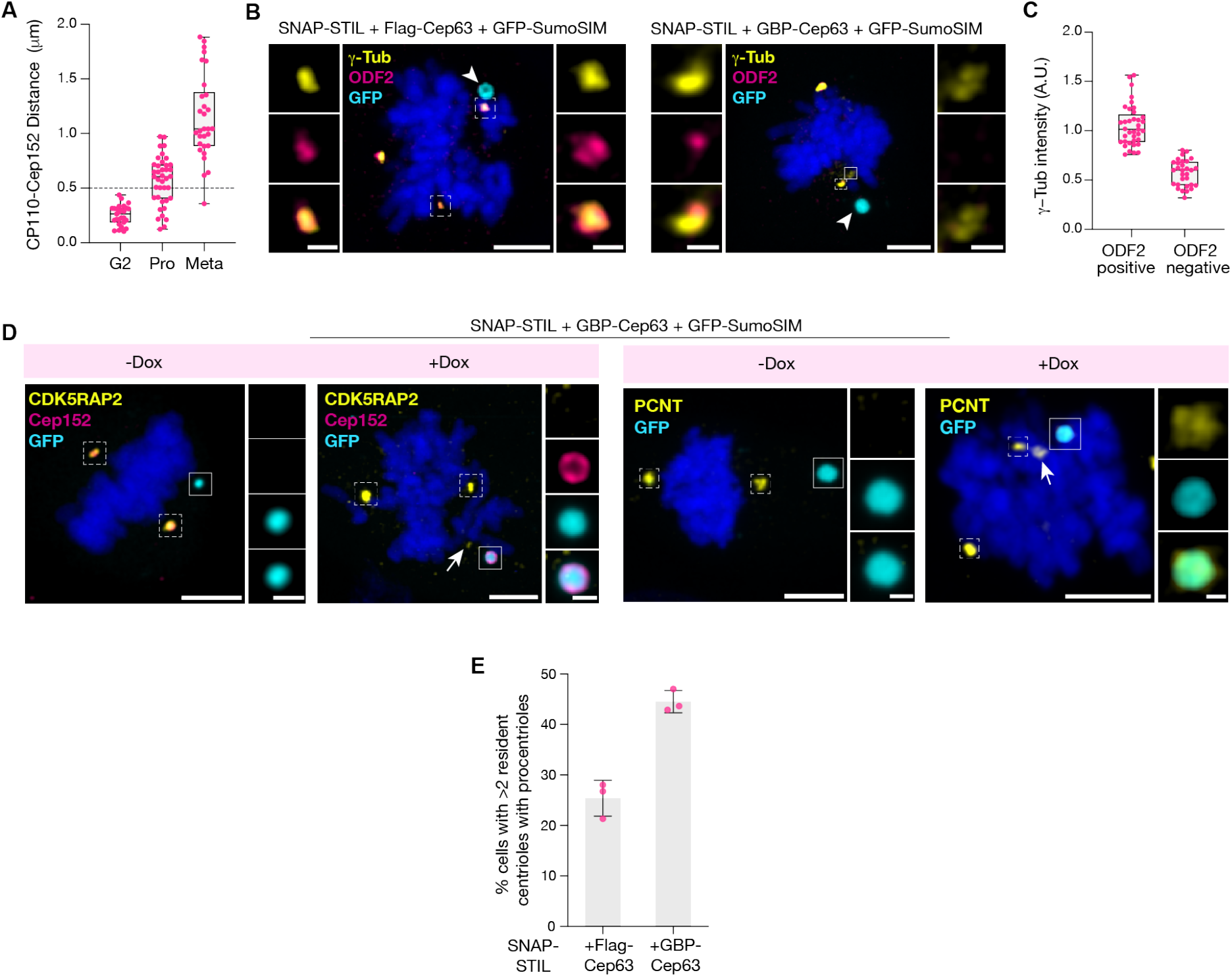
Droplet released ectopic procentrioles acquire PCM during mitosis. **A)** Quantification of distance between CP110 and Cep152 on droplet as shown in Fig 4A. Distances were measured from max projected images. n=25 for each condition. A cut-off distance of 0.5 μm, indicated by dashed line, was taken to consider procentrioles as released from the droplet. **B)** Representative images of mitotic cells expressing either Flag-Cep63 (left) or GBP-Cep63 (right), as well as SNAP-STIL, immunostained for γ-tubulin and ODF2. Arrowheads indicate droplet position. Dashed white boxes indicated γ-tubulin associated with ODF2 positive resident centrioles, solid white box γ-tubulin associated with ODF2 negative, droplet-released, procentrioles. **C)** Quantification of γ-tubulin intensity at ODF2 positive or ODF2 negative pole from the same cell. For each cell, intensity at the ODF2 positive pole is normalized to 1. Bars represent mean +/-s.d. n=30 cells were analyzed from three independent experiments. **D)** Representative images of mitotic cells immunostained for Cep152 and CDK5RAP2 or PCNT, as indicated. Arrow indicates weak CDK5RAP2 or PCNT pole close to the droplet targeted with Cep63. Images are representative examples from two experiments. Note that upon doxycycline addition, PCNT localizes weakly to Cep63 targeted droplets. **E)** Percentage of cells with more than two resident centrioles with associated procentrioles, as assayed by tubulin U-ExM. 50 cells, growing asynchronously, were counted at random for each condition from three independent experiments. Note that stable GFP-STIL over-expression in similar cells were reported to contain an even lower percentage of over-duplicating centrioles^67^. Scale bars: 5 μm in low magnification views, 1 μm in insets.

**Figure S7.**
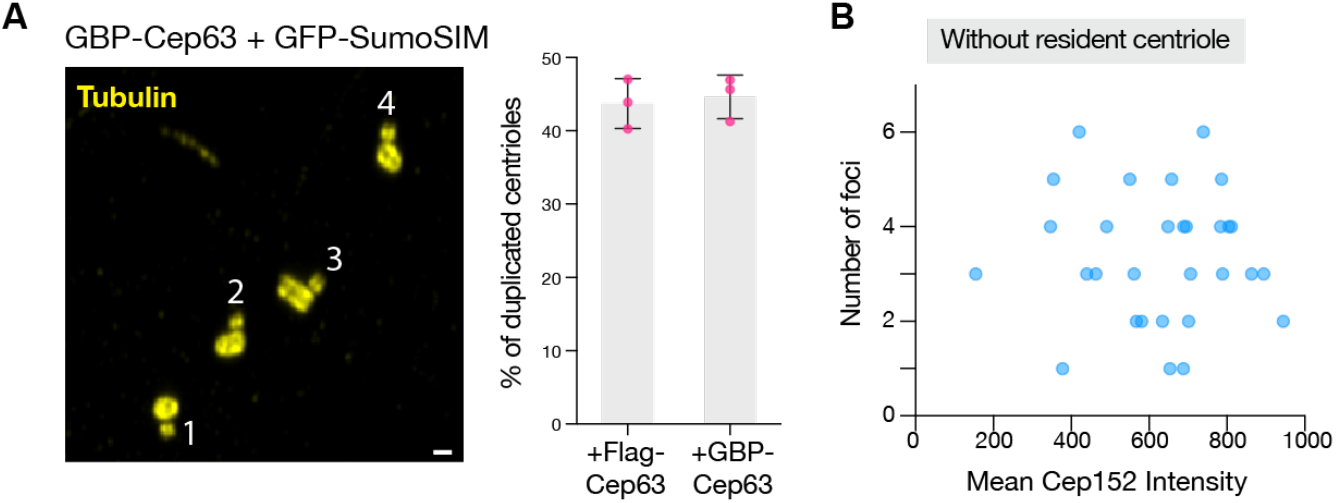
Procentrioles formed on the droplet surface in the absence of resident centrioles seed procentriole formation in turn. **A)** Left: representative tubulin U-ExM images of centrioles formed on the surface of the droplet in cells overexpressing SNAP-STIL and GBP-Cep63, 36 hours after Centrinone wash-out. Right: percentage of cells overexpressing SNAP-STIL and Flag-Cep63 or GBP-Cep63, in which resident centrioles have a neighboring procentriole 36 hours after Centrinone washout, as determined by tubulin U-ExM staining. 30 cells were counted for each condition from three independent experiments. Scale bar: 250nm. **B)** Mean Cep152 signal intensity on the droplet as a function of CP110 foci number Cep63 targeting to the droplet without resident centrioles. n=30 cells, each containing a single droplet were analyzed. Same droplets are reported in Figure 6G.

**Supplementary Movie 1 legend** U-2 OS cell expressing tagRFP-Centrin1 (red) together with Cep63-targeted droplet marked by GFP (cyan). Time zero is taken as the first instance of cell rounding indicating beginning of mitosis. Scale bar: 5 µm.

